# Combinations of *Spok* genes create multiple meiotic drivers in *Podospora*

**DOI:** 10.1101/562892

**Authors:** Aaron A. Vogan, S. Lorena Ament-Velásquez, Alexandra Granger-Farbos, Jesper Svedberg, Eric Bastiaans, Alfons J. M. Debets, Virginie Coustou, Hélène Yvanne, Corinne Clavé, Sven J. Saupe, Hanna Johannesson

**Author notes:** For correspondence: hanna. (HJ). These authors contributed equally to this work (Aaron A. Vogan, S. Lorena Ament-Velásquez).

## Abstract

Meiotic drive is the preferential transmission of a particular allele at a given locus during sexual reproduction. The phenomenon is observed as spore killing in a variety of fungal lineages, including *Podospora*. In natural populations of *Podospora anserina*, seven spore killers (*Psk*s) have been identified through classical genetic analyses. Here we show that the *Spok* gene family underlie the *Psk* spore killers. The combination of the various *Spok* genes at different chromosomal locations defines the spore killers and creates a killing hierarchy within the same population. We identify two novel *Spok* homologs that are located within a complex region (the *Spok* block) that reside in different chromosomal locations in given natural strains. We confirm that the individual SPOK proteins perform both the killing and resistance functions and show that these activities are dependent on distinct domains, a nuclease and a kinase domain respectively. Genomic data and phylogenetic analysis across ascomycetes suggest that the *Spok* genes disperse via cross-species transfer, and evolve by duplication and diversification within several lineages.

## Introduction

The genomes of all Eukaryotes harbour selfish genetic elements that employ a variety of mechanisms to undermine the canonical modes of DNA replication and meiosis to bias their own transmission (***Werren et al., 1988***; ***Burt and Trivers, 2009***). As the proliferation of these elements is independent from the regulated reproduction of the host organism, they can create conflict within the genome (***Rice and Holland, 1997***). Such intragenomic conflict is predicted by theory to spur an arms race between the genome and the elements, and consequently act as a major driver of evolutionary change (***Werren, 2011***). To understand the extent to which intragenomic conflict has shaped the evolution of genomes and populations it is crucial to identify the selfish genetic elements which are able to impact the dynamics of natural populations.

One important class of selfish genetic elements are known as meiotic drivers. These use a variety of mechanisms to hijack meiosis in order to bias their transmission to the gametes in proportions greater than 50% (***Sandler and Novitski, 1957***). This segregation distortion of alleles can be difficult to observe unless it is linked to an obvious phenotype such as sex (***Sandler and Novitski, 1957***; ***Helleu et al., 2014***), thus the prevalence of meiotic drive in nature is likely underestimated. Nevertheless, meiotic drive has been observed in many model systems, including *Drosophila, Mus, Neurospora*, and *Zea mays*, suggesting that it is likely widespread across all major Eukaryotic groups (***Lindholm et al., 2016***; ***Bravo Núñez et al., 2018***). Meiotic drive can be classified into three broad categories: female meiotic drive, sperm killing, and spore killing (***Lindholm et al., 2016***). Spore killing is found in ascomycete fungi and represents the most direct way to observe the presence of meiotic drive (***Turner and Perkins, 1991***). When a strain possessing a driving allele mates with a compatible strain that does not carry the allele (i.e., a sensitive strain), the meiotic products (ascospores) that carry the driving allele will induce the abortion of their sibling spores which do not have the allele. Spore killing is apparent in the sexual structures (asci) of the fungi as it results in half of the normal number of viable spores. Due to the haplontic life cycle of most fungi, spore killing is unusual among meiotic drivers as it is the only system where the offspring of an organism are killed by the drive (***Lyttle, 1991***). Additionally, with few exceptions (***Hammond et al., 2012***; ***Svedberg et al., 2018***), spore killer elements appear to be governed by single loci that confer both killing and resistance (***Grognet et al., 2014***; ***Nuckolls et al., 2017***; ***Hu et al., 2017***), which is in contrast to the other well-studied drive systems that comprise genomic regions as large as entire chromosomes (***Larracuente and Presgraves, 2012***; ***Hammer et al., 1989***).

Meiotic drivers are often expected to reach fixation or extinction in populations relatively rapidly (***Crow, 1991***), at which point the effects of the drivers will no longer be observable. In agreement with this expectation, most drivers which have been described exhibit large shifts in frequencies in both time and space (***Lindholm et al., 2016***; ***Carvalho and Vaz, 1999***). In the case of spore killers, multiple drivers have been found to coexist within a given species. The evolutionary dynamics of multiple drivers within species has not been thoroughly explored, but two contrasting examples are known. In genomes of *Schizosaccharomyces pombe*, numerous copies of both functional and pseudogenized versions of the *wtf* driver genes are found, suggesting that they duplicate readily, drive to high frequency in populations, and then lose their ability to kill (***Nuckolls et al., 2017***; ***Hu et al., 2017***). In contrast, the two spore killers *Sk-2* and *Sk-3* of *Neurospora intermedia* have only been described in wild strains four times and once respectively, whereas resistance to spore killing is widespread (***Turner, 2001***). In neither of these cases, have the impact of multiple drivers coexisting in a single population been characterized.

Natural populations of the filamentous fungus *Podospora anserina* are known to host multiple spore killers (***Grognet et al., 2014***; ***van der Gaag et al., 2000***; ***Hamann and Osiewacz, 2004***) and hence, provide an ideal system for the investigation of interactions among drivers at the population level. The first spore killer gene described in *P. anserina* was *het-s*, a gene that is also involved in allorecognition (***Dalstra et al., 2003***). Another class of spore killer genes in *Podospora* are known as *Spok* genes. *Spok1* is only known from a single representative of the closely related species *P. comata*, while *Spok2* has been shown to exist in high frequency among strains of a French population of *P. anserina* (***Grognet et al., 2014***). *Spok1* is capable of killing in the presence of *Spok2*, but not vice versa, indicating a dominant epistatic relationship between the two genes. Using visual observation of spore killing in crosses among French and Dutch *P. anserina* strains, seven separate spore killers have been identified (***van der Gaag et al., 2000***). These are referred to as *Psk-1* through *Psk-7* and can be distinguished through classical genetic analysis, by observing the presence, absence and frequency of killed spores when the different spore killers are crossed to each other (***Box 1***). At the onset of this study, it was not known whether the *Psk*s represent independent meiotic drive genes, or if they may be related to the *Spoks* and/or allorecognition loci. The *het-s* gene itself is not associated with the *Psk*s, but allorecognition is correlated with *Psk* spore killing (***van der Gaag et al., 2003***). On the other hand, the relationship between the *Spoks* is reminiscent of the hierarchy of killing among the *Psk*s, suggesting a possible connection between the activity of *Spok* genes and *Psk*s.

The primary goal of this study was to determine the identity of the genes that are responsible for the *Psk* spore killers found in *P. anserina*, and whether they relate to known meiotic drive genes. We identified two novel *Spok* homologs (*Spok3* and *Spok4*) and showed that these two, together with the previously described *Spok2*, represent the genetic basis of the *Psk* spore killers. The novel *Spoks* occur in large complex regions that can be found in different genomic locations in different strains. Our results illuminate the underlying genetics of a polymorphic meiotic drive system and expand our knowledge regarding their mechanism of action.

**Box 1 Figure 1.**
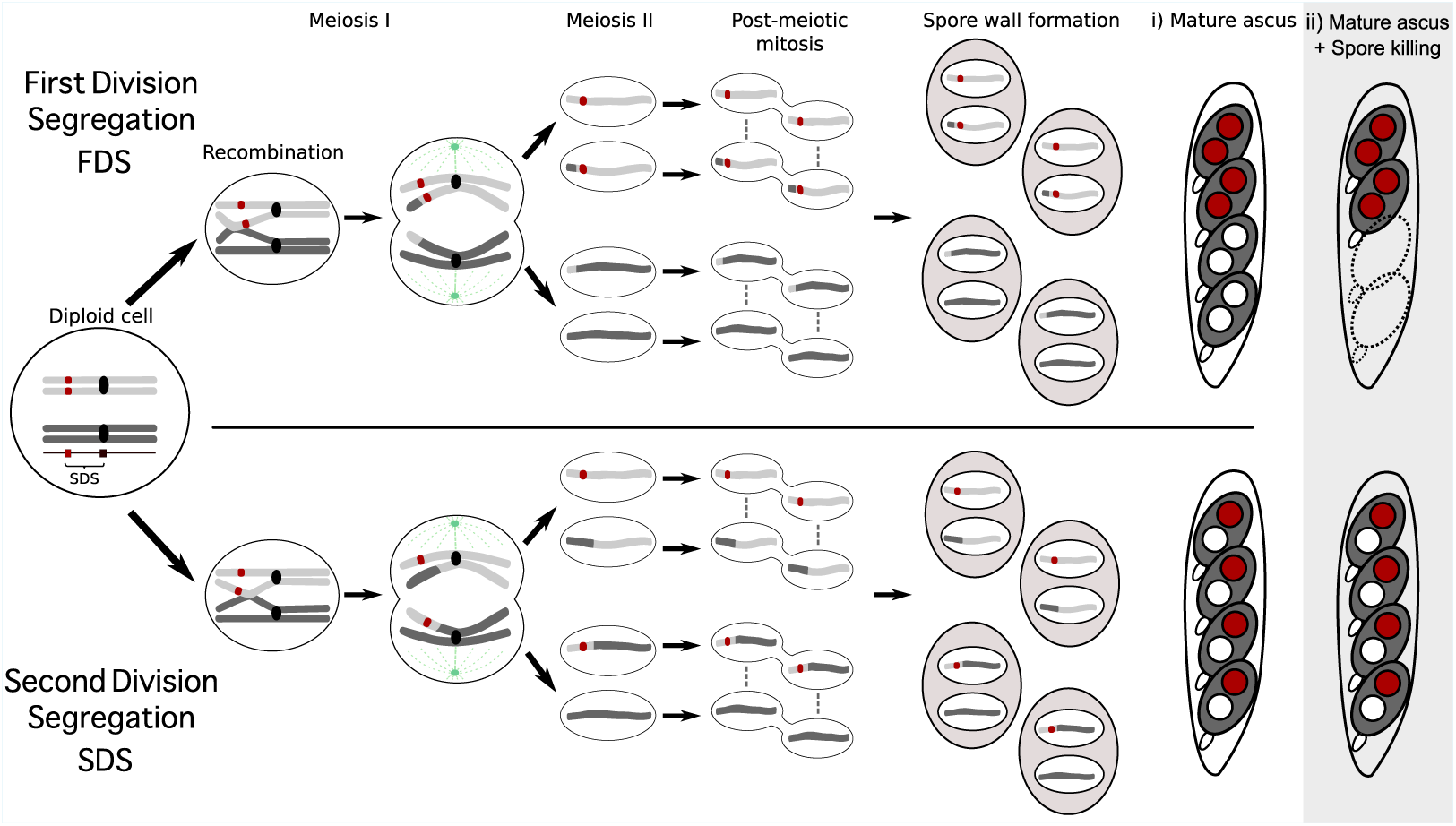
Meiosis and spore killing in *Podospora*. Meiosis in *Podospora* The seven separate *Psk*s are defined by their spore killing percentage and mutual interactions. To understand how the spore killing percentages relate to the genotypes of the strains, it is necessary to first appreciate some of the fundamental aspects of *Podospora* biology. Within the fruiting body (perithecium), dikaryotic cells undergo karyogamy to produce a diploid nucleus and immediately enter meiosis. After meiosis, one round of post-meiotic mitosis occurs, resulting in eight daughter nuclei. The nuclei are packaged together with their non-sister nuclei from mitosis (dashed line) to generate dikaryotic, self-compatible spores. In a cross where the parental strains harbour two alternative alleles for a given gene of interest (one of which is indicated by the red mark on the chromosome), spores can be produced which are either homoallelic or heteroallelic for the gene, depending on the type of segregation. Specifically, if there is no recombination event between the gene and the centromere, the gene undergoes first division segregation (FDS) and the parental alleles co-segregate during meiosis I, generating homoallelic spores (i). FDS of a spore killing gene will thus result in a 2-spored ascus (ii). If there is a recombination event between the gene of interest and the centromere, second division segregation (SDS) occurs. In this case heteroallelic spores will be formed (i). For spore killing, a 4-spored ascus will still be produced as only one copy of the spore killer is required to provide resistance (ii). As SDS is reliant on recombination, the frequency of SDS relates to the relative distance from the centromere and can be used for linkage mapping. When there is spore killing, the percent of 2-spored asci is the frequency of FDS, and is referred to as “spore killing percentage”. The *Psk*s were described by crossing different strains and evaluating what their spore killing percentage is in each cross. The seven unique *Psk*s were shown to interact in a complex hierarchy, showing either a dominance interaction, or mutual killing. Notably, crosses of strains carrying mutually resistant spore killers can still produce 2-spore asci if the killer loci are in different chromosomal locations (See ***Appendix 1*** for more details and ***Figure 4–Figure Supplement 1*** for a reproduction of the hierarchy presented in ***van der Gaag et al.*** (***2000***)).

## Results

### Genome assemblies

To investigate the genetic basis of spore killing in *P. anserina*, we generated high quality whole genome assemblies using a combination of long read (PacBio and MinION Oxford Nanopore) and short read (Illumina HiSeq) technologies. ***Table 1*** lists strains used for investigation. First, we selected strains from a natural population in Wageningen (Wa), the Netherlands, representing six of the previously described *Psk* spore killers (***van der Gaag et al., 2000***) along with a strain of a novel killing type (Wa100) that we referred to as *Psk-8*, and strain Wa63. Wa63 is of the same *Psk* type as the reference strain S, which we refer to herein as *Psk-S*. Additionally, we acquired and sequenced strains from the closely related *Podospora* species, *P. comata* (strain T) and *P. pauciseta* (CBS237.71). A strain labelled T (hereafter referred as T_G_) was kindly provided by Andrea Hamann and Heinz Osiewacz from the Goethe University Frankfurt and originates from the laboratory of Denise Marcou. However, as the genome sequence of T_G_ did not match that reported by ***Silar et al.*** (***2018***), but instead is a strain of *P. anserina*, we included in our dataset another strain labelled T from the Wageningen Collection that was originally provided by the laboratory of Léon Belcour. We refer to this strain as T_D_, and sequenced it using only Illumina HiSeq. The genome of T_D_ matches ***Silar et al.*** (***2018***) as the epitype of *P. comata* (See ***Appendix 2*** for further discussion).

**Table 1.**
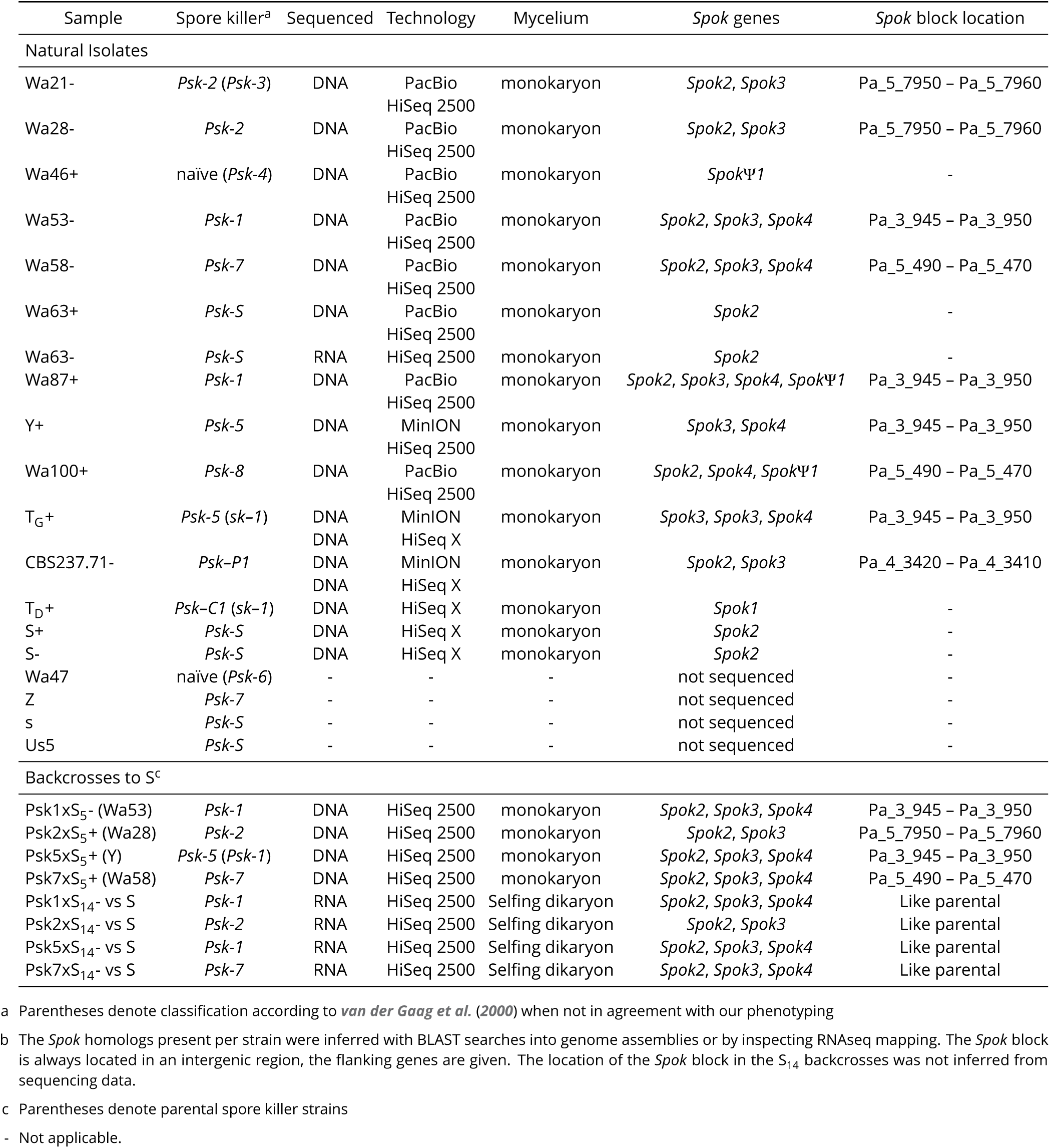
List of all strains used in this study.

The final assemblies (long-read technologies polished with Illumina HiSeq data) consist of 18 to 53 scaffolds, from which the majority were either mitochondrial or rDNA in origin. Amongst the remaining scaffolds, the expected seven chromosomes were recovered in their entirety for almost all strains with PacBio data, and in up to 15 scaffolds with those sequenced using MinION (***Figure 1–source data 1***). Since the assemblies of each strain were produced from one haploid (monokaryotic) isolate, we will refer to specific genome assemblies with their strain name followed by their corresponding mating type, e.g. Wa63+.

**Figure 1.**
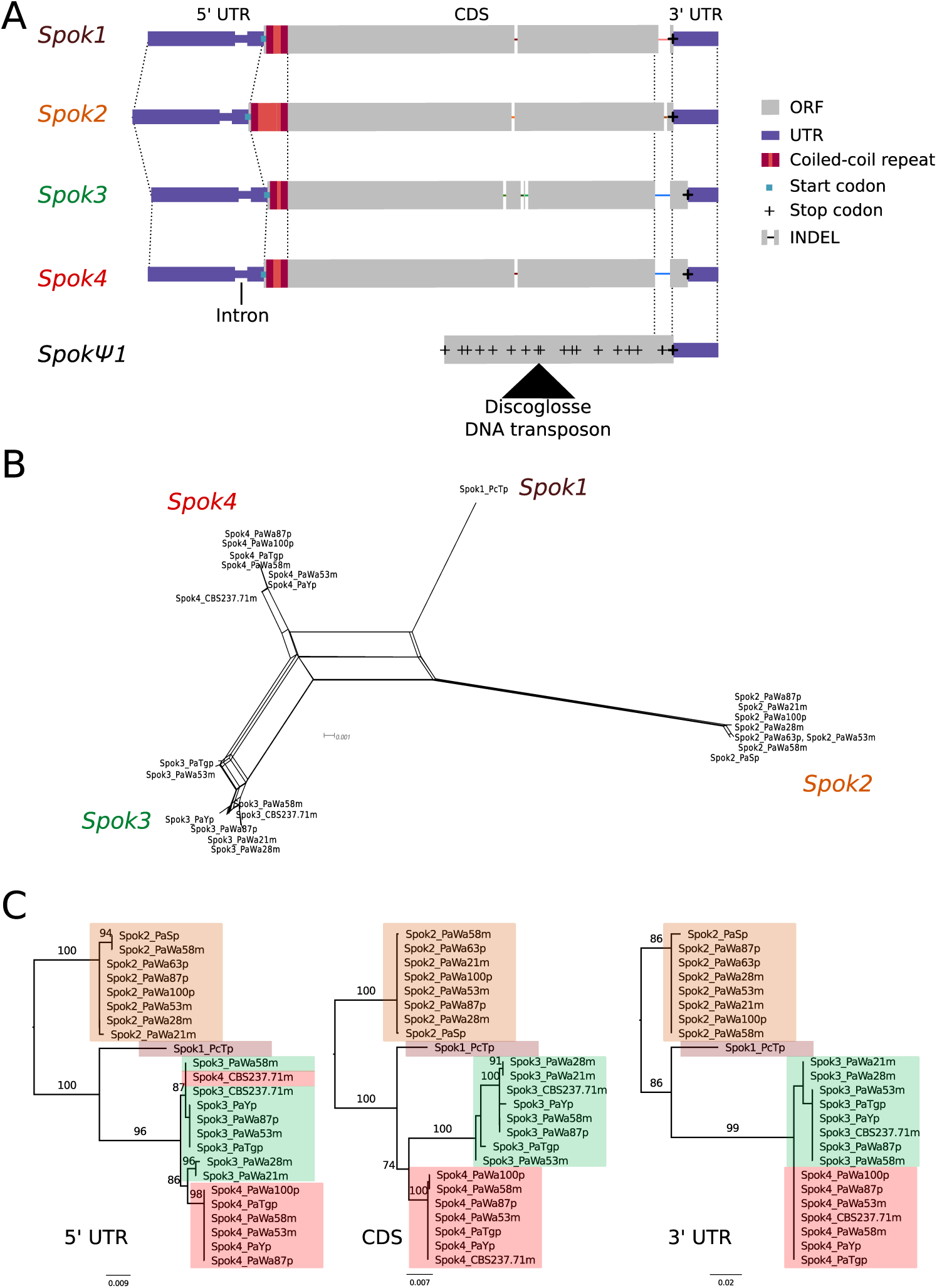
Relationships among the *Spok* homologs. **A** Schematic representation of the main features of the *Spok* genes. All homologs share an intron within the 5’ UTR. At the start of the coding region there is a repeat region, where the number of repeats varies among the homologs. The central portion of the coding regions has a number of indels, which appear to be independent deletions in each of *Spok2, Spok3*, and *Spok4*. There is a frameshift mutation at the 3’ end of the coding region that shifts the stop codon of *Spok3* and *Spok4* into what is the 3’ UTR of *Spok1* and *Spok2*. The pseudogenized *Spok* gene contains none of the aforementioned central indels and appears to share the stop codon of *Spok1* and *Spok2*. However, there are numerous mutations resulting in stop codons within the CDS as well as a full DNA transposon (discoglosse). No homologous sequence of the 5’ end of the pseudospok is present. **B** A NeighborNet split network of all active *Spok* genes from all strains sequenced in this study. The four homologs cluster together well, however there are a number of reticulations, presumably due to gene conversion events. **C** Maximum likelihood trees based of three separate regions of the *Spok* genes: the 5’ UTR, the CDS, and the 3’ UTR (starting from the stop codon of *Spok3* and *Spok4*). The trees are rooted arbitrarily using *Spok2*. Branches are drawn proportional to the scale bar (substitutions per site), with bootstrap support values higher than 70 shown above. **Figure 1–Figure supplement 1.** Visualised nucleotide alignment. **Figure 1–Figure supplement 2.** *Spok* transcripts. **Figure 1–source data 1.** Nucleotide alignment of *Spok* genes. **Figure 1–source data 2.** Splits tree in Nexus format.

### Identification of novel *Spok* genes

By searching our assemblies for the *Spok2* sequence (presented by ***Grognet et al.*** (***2014***)) using BLAST, we could confirm the presence of this *Spok* gene in the majority of strains, in agreement with ***Grognet et al.*** (***2014***). Furthermore, based on sequence similarity with *Spok2*, we identified two novel homologs that we refer to as *Spok3* and *Spok4*. Additionally, the BLAST searches recovered a pseudogenized *Spok* gene (*Spok*Ψ*1*). The *Spok* gene content of the strains investigated in this study is reported in ***Table 1***.

A schematic representation of the *Spok* homologs is shown in ***Figure 1 A***. We considered the *Spok2* sequence of S+, and the *Spok3* and *Spok4* sequences of Wa87+ as reference alleles for each homolog. Overall they show a high degree of conservation, including the 3’ and 5’ UTRs. A nucleotide alignment of the *Spok* genes’ CDS revealed 130/2334 variable sites among the homologs (***Figure 1–Figure Supplement 1*** and ***Figure 1–source data 1***). A relatively large proportion (67%, 87/130) of those result in amino acid changes and 74% are unique to one of the *Spok* homologs. There are also six indels among all the *Spok* genes including one at the 5’ end of the ORF, which represents a variable length repeat region, and one at the 3’ end of the ORF shared by *Spok3* and *Spok4*. The 3’ end indel induces a frameshift and changes the position of the stop codon (***Figure 1A***). *Spok*Ψ*1* has a missing 5’ end, multiple stop codons, and a discoglosse (Tc1/*mariner*-like) DNA transposon (***Espagne et al., 2008***) inserted in the coding region. Of particular interest, *Spok*Ψ*1* has no deletions relative to the other *Spok* homologs, suggesting the indels in the functional *Spok* homologs represent derived deletions.

To aid in the identification of the meiotic drive genes, we gathered Illumina HiSeq data from the reference strain S together with four strains resulting from backcrossing of *Psk-1, Psk-2, Psk-5*, and *Psk-7* into S (***Table 1***). These and all other genomes sequenced with short-read data were assembled de novo using SPAdes. The resulting assemblies consisted of between 222 and 418 scaffolds larger than 500bp, with a mean N50 of 227 kbp (Supplementary file 2). The coverage of the publicly available reference genome of the strain S+ (***Espagne et al., 2008***), hereafter referred to as Podan2, was above 98% for all of the SPAdes assemblies of *P. anserina*. When the filtered Illumina reads were mapped to Podan2, all samples had a depth of coverage above 75x (Supplementary file 2). Taken together, our genome assemblies, resulting from both long and short-read data, are very comprehensive.

There is little allelic variation within the *Spok* homologs in the Wageningen population and the variants of the four homologs cluster phylogenetically (***Figure 1B and C***). The *Spok2* gene in the Wageningen strains are identical to the two alleles described in ***Grognet et al.*** (***2014***), with the exception of *Spok2* from Wa58-which has a single SNP that results in a D358N substitution. The *Spok2* allele of the French strain A, which shows resistance without killing (as reported by ***Grognet et al.*** (***2014***)), was not found in any of our genomes. *Spok3* has five allelic variants, and the allelic variation of *Spok4* is reminiscent of *Spok2* with only Wa100+ and Wa58-having a single synonymous SNP (***Figure 1*C**). Lastly, the three copies of *Spok*Ψ*1* are all unique (***Figure 2–source data 2***).

**Figure 2.**
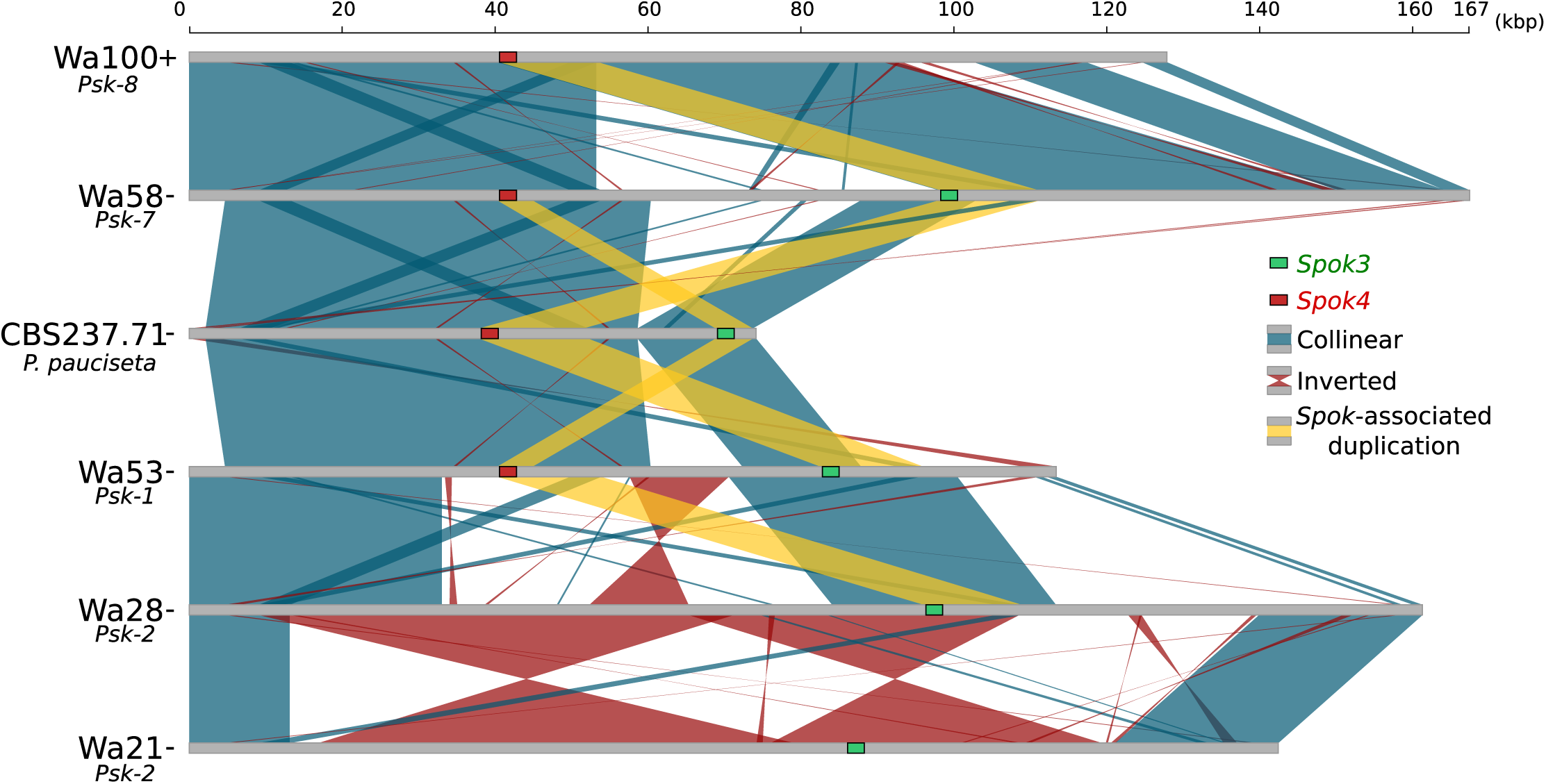
Alignment of the *Spok* blocks from different strains. Grey bars represent the block sequences, blue vertical lines connect collinear regions between blocks, while red lines indicate inverted regions. The yellow lines show the region that is duplicated within the block surrounding *Spok3* (green) and *Spok4* (red). **Figure 2–Figure supplement 1.** Alignment of *Psk-1/5 Spok* blocks. **Figure 2–Figure supplement 2.** Dot plot showing sytneny between the *Spok*Ψ*1* region and the *Spok* block. **Figure 2–Figure supplement 3.** Introgressed regions of the S_5_ backcrossed strains. **Figure 2–source data 1.** Fasta File of the *Spok* block from all strains. **Figure 2–source data 2.** Fasta File of the *Spok*Ψ*1* region from all strains.

Notably, a number of the variants of *Spok3* show signatures of gene conversion events (***Lazzaro and Clark, 2001***). Specifically, strain Y+ has three SNPs near the start of the gene that result in amino acid changes and match exactly those in *Spok2* (***Figure 1–Figure Supplement 1***). The Wa53+ allele of *Spok3* has a series of SNPs (a track of 205 bp) that are identical to *Spok4*, but different from all other *Spok3* sequences, and three additional SNPs near the 5’ end that also match *Spok4* (***Figure 1–Figure Supplement 1***). The T_G_+ strain possesses two identical copies of *Spok3* (see **Methods**) that share the aforementioned tract with Wa53+, but which extends for an additional 217 bp (***Figure 1–Figure Supplement 1***). These chimeric *Spoks* are recovered from the final assemblies (pre-and post-Pilon polishing) with high long-read coverage (>30x), suggesting that our finding is not a bioinformatic artifact. The gene conversion events between *Spok* homologs are supported by the reticulation shown in a NeighborNet split network (***Figure 1*B**) and by a significant recombination Phi test (199 informative sites, p = 1.528e-12). A Maximum Likelihood phylogenetic analysis of the UTR sequences (defined by conservation across homologs) suggests that *Spok3* and *Spok4* are closely related (***Figure 1*C**), which is at odds with the high structural similarity of the CDS of *Spok1* and *Spok4* (***Figure 1*A**). Therefore, we cannot make any strong inference about the relationships between the *Spok* homologs from the sequence data.

The *Spok1* gene was previously identified from T_D_ (***Grognet et al., 2014***). No other strains investigated in this study were found to possess *Spok1*, indicating that it is likely not present in *P. anserina*. Remarkably, BLAST searches of the *Spok2* with the UTR sequences revealed the presence of a small piece (∼156 bp long) of a presumably degraded *Spok* gene in the T_D_ de novo assembly and on the chromosome 4 of the reference *P. comata* genome released by ***Silar et al.*** (***2018***). This piece overlaps with the last amino acids of the CDS 3’ end and it is flanked by an arthroleptis (solo LTR) retrotransposon on one side and by unknown sequence on the other. Due to the small size, it is unclear if this piece belongs to a novel *Spok* gene, but the location (between genes PODCO_401390 and PODCO_401400) does not align with any other known homolog. Strain CBS237.71 was formerly identified as *P. comata* and was reported to possess a *Spok* gene (***Grognet et al., 2014***). It has now been assigned to its own species, *P. pauciseta* (***Boucher et al., 2017***) and the sequencing reveals that the genome of this strain contains both *Spok3* and *Spok4* (***Figure 1*B**).

### Backcrossing confirms the association of the *Spok* genes with the *Psk***s**

Four of the *Psk* spore killers were previously introgressed into the reference strain S through five successive backcrosses (***van der Gaag et al., 2000***) and are referred to here as Psk1xS_5_, Psk2xS_5_, Psk5xS_5_, and Psk7xS_5_ (***Table 1***). Our Illumina data recovered in total 41482 filtered biallelic SNPs from the four S_5_ backcrosses and the parental strains. All backcrossed strains show a few continuous tracts of SNPs from the killer parent (***Figure 2–Figure Supplement 3***). For example, Psk1xS_5_-has a long tract in chromosome 1 that represents the mat-mating type, which is expected since the published reference of S (Podan2), for which the SNPs are called, is of the opposite mating type (mat+). Importantly, the location of the *Spok* genes of each parental strain has a corresponding introgressed SNP tract in its S_5_ backcross, while all backcrossed strains possess the *Spok2* gene from strain S (***Figure 2–Figure Supplement 3***). Notably, crossing results reveal that Psk5xS_5_ has a *Psk-1* killing phenotype whereas all other S_5_ backcrossed strains maintained the parental phenotype (***Figure 4–source data 1***). However as strain Y does not possess *Spok2*, the overall *Spok* content of Y is not the same as Psk5xS_5_ (***Table 1***). These data suggest that the *Spok* content is responsible for the killer phenotype of the *Psk*s.

As the various *Psk* types reflect specific *Spok* gene content, we can estimate the frequency of each *Spok* gene in the Wageningen population from ***van der Gaag et al.*** (***2000***). We have determined the *Spok* gene composition for *Psk-1, Psk-2, Psk-4, Psk-5*, and *Psk-7*, as well as those previously considered as “sensitive”, now *Psk-S*. These account for 92/99 strains collected from Wageningen. The seven remaining strains were identified as either *Psk-3* or *Psk-6*. Our representative strain of *Psk-3* (Wa21) was shown to be *Psk-2*, and we are unable to comment on *Psk-6* as our representative strain (Wa47) behaves as *Psk-4* in test crosses (***Table 1***). Therefore we assume strains annotated as *Psk-4* possess no functional *Spok* genes (hereafter referred to as naïve) and omit all the *Psk-3* strains (except Wa21) and the *Psk-6* strains (except Wa47) from the analysis. Hence, *Spok2* is estimated to be in 98% of strains, *Spok3* in 17%, and *Spok4* in 11% of Dutch strains.

### *Spok* genes are found in complex regions associated with killer phenotypes

While the *Spok* genes are often assembled into small fragmented contigs when obtained by using Illumina data alone, in the PacBio and MinION assemblies *Spok3* and *Spok4* are fully recovered within an inserted block of novel sequence (74–167 kbp depending on the strain), hereafter referred to as the *Spok* block. When present, the *Spok* block was never found more than once per genome and always contains at least one *Spok* gene. Whole genome alignments revealed that the *Spok* block has clear boundaries, and is localized at different chromosomal positions on chromosome 3 or in either arm of chromosome 5 in different strains of *P. anserina* (***Table 1***). Importantly, these positions correspond with a single SNP tract from the S_5_ backcrosses. In *P. pauciseta* (CBS237.71) the *Spok* block is found in chromosome 4. The *Spok* block of the different strains shares segments and overall structure (***Figure 2*** and ***Figure 2–Figure Supplement 1***), which suggests that they have a shared ancestry. However, complex rearrangements are found when aligning the block between the genomes. Within the *Spok* block, a given strain can harbour either or both of *Spok3* and *Spok4* and the regions containing the *Spok* genes appear to represent a duplication event (***Figure 2***). Strain T_G_+ shows an additional duplication which has resulted in a second copy of *Spok3* (***Figure 2–Figure Supplement 1***). While *Spok3* and *Spok4* are always found within the block, *Spok2* is never associated with a *Spok* block, but is found at the same location on chromosome 5 as previously described for the reference strain S (***Grognet et al., 2014***). When present, *Spok*Ψ*1* was found at a single position in the right arm of chromosome 5. It is surrounded by numerous transposable elements (TEs), and the region does not appear to be homologous to the *Spok* block (***Figure 2–Figure Supplement 2***).

In the few strains with no copy of *Spok2*, analysis of the region suggests that this is a result of a one-time deletion (***Figure 3***). The annotation in the original reference genomes of T_D_ and S is erroneous due to misassemblies and/or incomplete exon prediction, which were both corrected using our own Illumina data, annotation pipeline, and validated with RNAseq expression data of T_D_. First, the flanking gene P_5_20 (marked as (1) in ***Figure 3***) in *P. pauciseta* (CBS237.71) and *P. comata* (T_D_) is considerably longer than the *P. anserina* ortholog, which is truncated by a discoglosse (Tc1/*mariner*-like) DNA transposon (2). In the strains without *Spok2* (Wa46, Y, and T_G_), this discoglosse itself is interrupted and the sequence continues on the 3’ end of a fragmented crapaud (*gypsy*/Ty3) LTR element, which can be found in full length downstream of *Spok2* in the other strains. This configuration implies that the absence of *Spok2* constitutes a deletion (3), rather than the ancestral state within *P. anserina*. An alternative scenario would require multiple additional insertions and deletions of TEs and *Spok2*.

**Figure 3.**
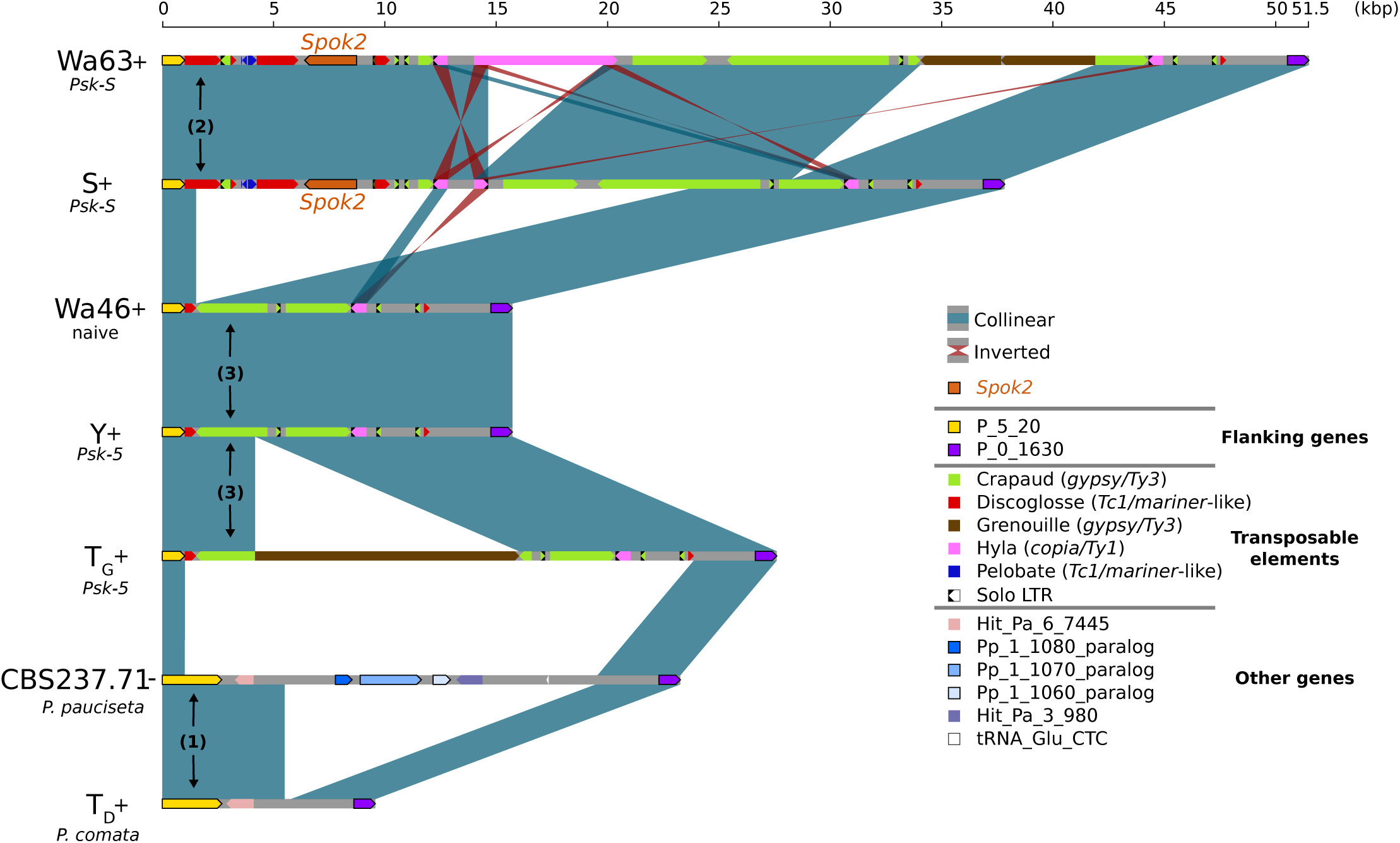
Alignment of the *Spok2* locus in selected strains. The haplotypes are defined by the flanking genes P_5_20 and P_0_1630 located in chromosome 5 of the three sampled species. Every strain has a haplotype of different size, mainly due to differences in transposable element (TE) content. Within *P. anserina*, the TE variation across all sequenced strains occurs downstream of *Spok2*, as exemplified by strains Wa63 and S. The strains Wa46, Y and T_G_ all lack *Spok2* and share break points. Notice that P_5_20 stands for the Pa_5_20 and PODCO_500020 in the reference annotation of *P. anserina* and *P. comata*, respectively, while P_0_1630 stands for Pa_0_1630 and PODCO_001630. As a note, *P. pauciseta* has a duplication of three genes in tandem from chromosome one (Pa_1_1080-60) between the flanking genes. Hit_Pa_X_XXX genes stand for significant BLAST hits to genes of Podan2. TE nomenclature follows ***Espagne et al.*** (***2008***). **Figure 3–source data 1.** Fasta File of the *Spok2* region from all strains. **Figure 3–source data 2.** Annotation File for TEs surrounding *Spok2*.

### *Spok3* and *Spok4* function as meiotic drive genes

We constructed knock-in and knock-out strains to confirm that the newly discovered *Spok* homologs *Spok3* and *Spok4* can induce spore killing on their own (***Table 2***), as previously shown for *Spok2* by ***Grognet et al.*** (***2014***). First, the *Spok2* gene was deleted from the strain s to create a Δ *Spok2* strain for use with the knock-ins. A cross between s and the Δ *Spok2* strain resulted in about ∼40%of 2-spored asci as previously reported by ***Grognet et al.*** (***2014***), (80/197, 40.6%) (***Figure 4–Figure Supplement 2B***). The *Spok3* and *Spok4* genes were inserted separately at the centromere-linked *PaPKS1* locus (a gene controlling pigmentation of spores (***Coppin and Silar, 2007***)). A *Spok3::PaPKS1* Δ *Spok2* x Δ *Spok2* cross yielded almost 100% 2-spored asci with two white (unpigmented) spores (118/119, 99.1%) (***Figure 4– Figure Supplement 2C***). Similarly, a *Spok4::PaPKS1* Δ*Spok2* x Δ*Spok2* cross yielded almost 100% 2-spored asci with two white (unpigmented) spores (343/346, 99.1%) (***Figure 4–Figure Supplement 2D***), indicating that *Spok3* and *Spok4* function as spore killers when introduced in a single copy at the *PaPKS1* locus.

**Table 2.**
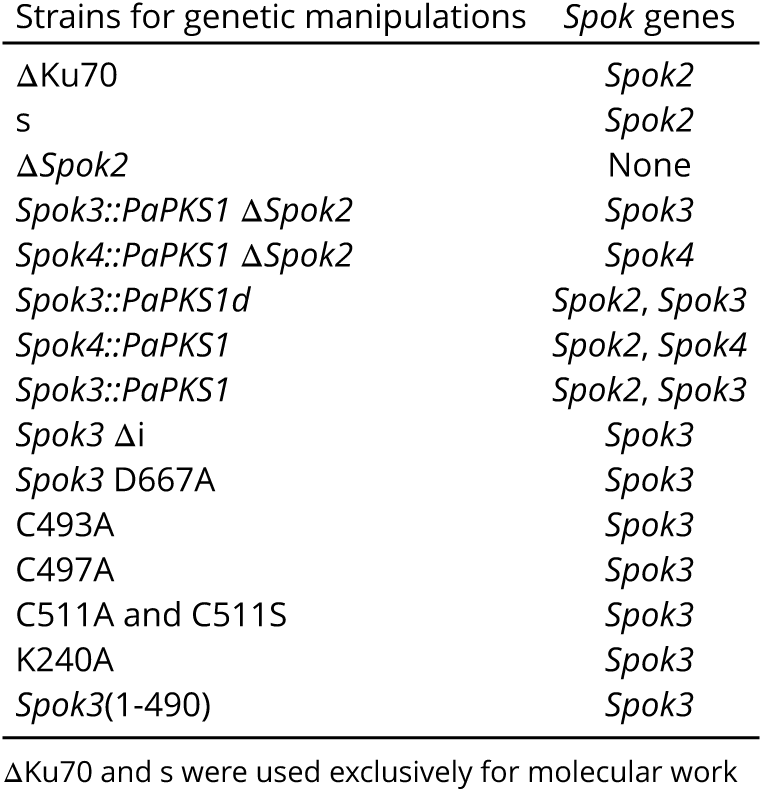
*Spok* gene content of genetically modified strains.

**Figure 4.**
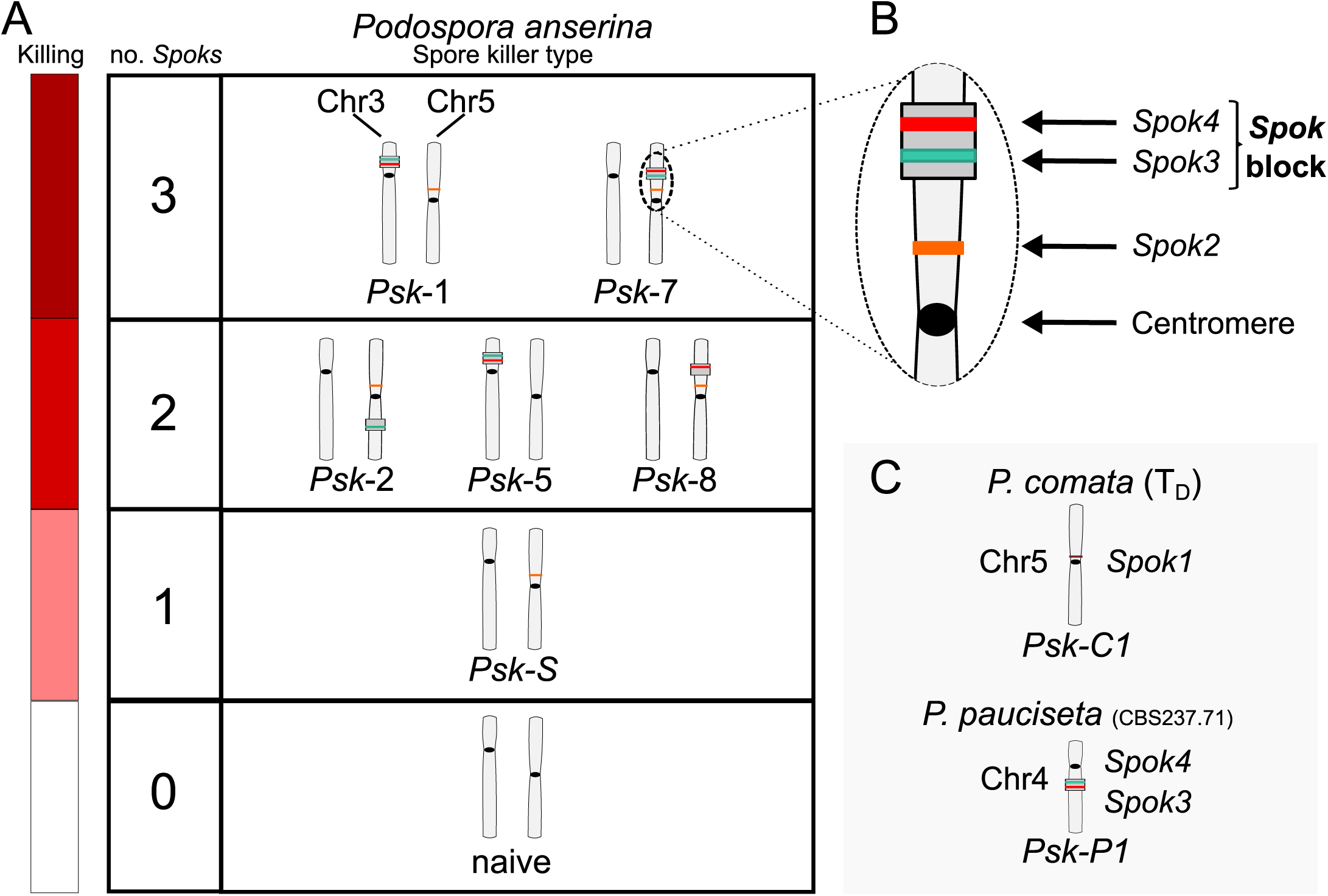
Interactions among the various *Psk* types and the occurrence of *Spok* genes. **A** The boxes represent hierarchical levels that increase in killing dominance from bottom to top, which correlates with the number of *Spok* genes that a strain possesses. Strains with three *Spok* genes induce spore killing of strains with only two *Spok* genes and show mutual resistance to each other. Strains with two *Spok* genes show mutual killing among themselves due to the different *Spok* genes and kill strains with only *Spok2*. Strains with one *Spok* kill strains with no *Spok* genes (naïve strains). The chromosome diagrams depict the presence of the *Spok* genes and their location in the genome for the sequenced strains. **B** A zoomed in look at Chromosome 5 of a *Psk-7* strain demonstrating that *Spok3* and *Spok4* are present in the *Spok* block and *Spok2* is present at the standard location. **C** The closely related species *P. comata* and *P. pauciseta* also possess *Spok* genes, but at different locations. The *Spok* genes in *P. pauciseta* are present in a smaller *Spok* block, while *Spok1* is found on its own and exclusively in *P. comata*. **Figure 4–Figure supplement 1.** Depiction of *Psk* killing heirarchy from ***van der Gaag et al.*** (***2000***). **Figure 4–Figure supplement 2.** Images of spore killing between genetically modified strains. **Figure 4–Figure supplement 3.** Results from pooled sequencing experiment of a cross between *Psk-1* and *Psk-5*. **Figure 4–source data 1.** Table with killing percentages for all crosses tested between strains. **Figure 4–source data 2.** Table with killing percentages for test crosses to determine epistatic interactions.

### The *P. anserina Spok* homologs are functionally independent

To determine whether there are epistatic interactions among the *Spok* genes of *P. anserina*, pairwise crosses between the strains were conducted to determine which matings resulted in spore killing (***Figure 4–source data 1***). To assess any epistatic interaction between different killer types, dikaryotic F1 progeny that are homoallelic for the killing locus (***Box 1***) were selected, backcrossed to both parental strains, and were also allowed to self. Killing interactions were classified into one of the following categories.

1. Dominance interaction - Spore killing is observed when backcrossed to only one of the parental strains, and no spore killing is observed upon selfing.
2. Mutual resistance - No spore killing is observed when backcrossed to either parent nor when selfed.
3. Mutual killing - Spore killing is observed when backcrossed to either parent and/or when F1 progeny are selfed.

As an example, in a cross between *Psk-1* and *Psk-7* there is spore killing. However the F1 progeny from this cross show no killing to either parent, satisfying condition 2. Thus they are mutually resistant, which is consistent with the fact that they carry the same three *Spok* genes. The reason spore killing is observed in the original cross is because the *Spok* block is located at different genomic positions. As a result, the *Spok* block can co-segregate during meiosis, leaving two spores without any *Spoks* and making them vulnerable to killing (see ***Appendix 1*** for a detailed explanation).

The results from these crosses are reported in ***Figure 4–source data 1***. Note that we found several of the *Psk* designations of the strains to differ from those reported previously, and these discrepancies are shown in ***Table 1***. From the epistatic interactions and killing percentages of the crosses, we construct a killing hierarchy (***Figure 4***) that also differs from that reported in ***van der Gaag et al.*** (***2000***) (***Figure 4–Figure Supplement 1***). In summary, our results show that *Spok2, Spok3*, and *Spok4* all act as spore killers and have no epistatic interactions with each other. The killing hierarchy observed in the Wageningen population of *P. anserina* is an emergent property of the presence and absence of the various *Spok* homologs in the different genomes. The rationale behind these conclusions is explained in detail below.

Spore killer types *Psk-1* and *Psk-7* reside at the top of the hierarchy, possess a *Spok* block with both *Spok3* and *Spok4*, and have *Spok2* (***Figure 4***). *Psk-2* and *Psk-8* are both dominant over *Psk-S*, which only has *Spok2*. *Psk-2* has a *Spok* block with just *Spok3* on the right arm of chromosome 5 and *Psk-8* has a *Spok* block with just *Spok4* at the same position as *Psk-7* on the left arm of chromosome 5, indicating that *Spok2* does not provide resistance to either *Spok3* or *Spok4*. *Psk-1* and *Psk-7* are both dominant over *Psk-2* and *Psk-8*, indicating that *Spok3* does not provide resistance to *Spok4* and vice versa. The fact that *Psk-S* is capable of killing strains with no *Spok* genes (i.e. naïve) confirms previous results that *Spok2* alone is able to induce spore killing (***Figure 4***; and see ***Grognet et al.*** (***2014***)).

*Psk-5* is a slightly more complicated case. It displays mutual killing with *Psk-S* and kills naïve strains, but *Psk-1* is dominant over *Psk-5*. *Psk-1* and *Psk-5* possess the same *Spok* block at the same location in Chromosome 3 (***Figure 4*** and ***Figure 2–Figure Supplement 1***), but *Psk-5* does not possess *Spok2*, suggesting that *Spok2* is responsible for killing in these crosses. If *Spok2* is responsible for killing when *Psk-1* is crossed with *Psk-5*, we expect the killing percentage to be the same as with crosses between *Psk-S* and naïve strains (∼40%). However, these crosses consistently show only ∼25% killing. To confirm that *Spok2* is responsible for killing in crosses between *Psk-1* and *Psk-5*, a pooled sequencing approach was employed. A cross was conducted between Wa87 (*Psk-1*) and Y (*Psk-5*), and spores from 2-spored (spore killing) and 4-spored asci (heteroallelic for killers) were collected and sequenced in separate pools. The 2-spored pool only contains SNPs from Wa87 for a large portion of Chromosome 5, which includes the *Spok2* gene, whereas the 4-spored pool contains SNPs from both parents at this genomic location (***Figure 4–Figure Supplement 3***). As the 2-spored asci are the result of FDS of the killing locus (***Box 1***), this result strongly suggests that *Spok2* is responsible for spore killing when *Psk-1* is crossed to *Psk-5* and thus that neither *Spok3* nor *Spok4* provides resistance against *Spok2*.

Of note, crosses between *Psk-1* and *Psk-5* often produce 3-spored asci and occasionally show erratic killing, which may contribute to the lower killing percentages. This phenomenon is also observed in crosses between *Psk-S* and naïve strains. We have been able to isolate a spore from a 3-spored ascus in a cross between *Psk-S* and a naïve strain that has no copy of *Spok2* (***Appendix 2***). Therefore, the 3-spored asci are likely due to incomplete penetrance of the killing factor and supports the conclusion that the spore killing observed in these crosses is caused by the same gene, *Spok2*. This result is consistent with findings presented in the study by ***van der Gaag*** (***2005***) that provided independent evidence for incomplete penetrance of spore killing between S and Wa46 (*Psk-S* and naïve).

The spore killing interactions of *Spok3* and *Spok4* cannot be dissociated from the *Spok* block with the use of wild or introgressed strains, so we made use of the aforementioned knock-in strains to confirm the independence of the *Spok* gene interactions from the *Spok* block. First, to confirm the killing interaction between *Spok3* and *Spok4*, we crossed a strain bearing *Spok4* at *PaPKS1* with a strain bearing *Spok3*. Because crosses homozygous for the *PaPKS1* deletion have poor fertility, we constructed a strain in which *Spok3* is inserted as a single copy at the *PaPKS1* locus but just downstream of the coding region (*Spok3::PaPKS1d*) in order to yield strains with normal pigmentation and normal fertility in crosses to *PaPKS1* deletion strains. In control crosses, the *Spok3::PaPKS1d* strain showed killing when crossed with a strain lacking *Spok3* but no killing when crossed with *Spok3::PaPKS1* (***Figure 4–Figure Supplement 2E and F***). The cross between *Spok3::PaPKS1d* and *Spok4::PaPKS1* yields asci with 4 aborted spores indicating mutual killing of *Spok3* and *Spok4* (***Figure 4–Figure Supplement 2G***). To determine the killing relation between *Spok2* and *Spok3*, a cross was conducted between *Spok3::PaPKS1* and s. This cross yielded mostly 2-spored asci with two unpigmented spores (163/165, 98.8%) (***Figure 4–Figure Supplement 2H***) indicating that *Spok3* kills in the presence of *Spok2*. Similarly, to determine the killing relation between *Spok2* and *Spok4*, a cross was conducted between *Spok4::PaPKS1* and s (216/217, 99.5%) (***Figure 4–Figure Supplement 2I***). While these crosses indicate that *Spok2* does not confer resistance to *Spok3* and *Spok4* (*Spok3* and *Spok4* both kill *Spok2*), they do not allow us to determine as such whether *Spok3* or *Spok4* confer resistance to *Spok2*. To address this point, *Spok2* killing was analyzed in a cross homozygous for *Spok3* (*Spok3::PaPKS1* x *Spok3::PaPKS1d* Δ*Spok2*), which yielded 46% two-spored asci (143/310) confirming that *Spok2* killing occurs in the presence of *Spok3* (***Figure 4–Figure Supplement 2J***). To determine if *Spok4* is resistant to *Spok2*, we made a *Spok4::PaPKS1* x *Spok4::PaPKS1* Δ*Spok2* cross (11/24 two-spored asci) (***Figure 4–Figure Supplement 2K***). Although this genetic background is ill suited for determining killing frequency (because of the aforementioned effect of the homozygous *PaPKS1* deletion on fertility), presence of 2-spore asci suggests that *Spok4* does not confer resistance to *Spok2* killing. Overall, these results confirm the findings with the wild strains that *Spok2, Spok3*, and *Spok4* have no epistatic interactions, and imply that the *Spok* block does not augment the function of the *Spok* genes.

In contrast to the absence of epistatic interactions among *Spok* genes of *P. anserina, Spok1* of *P. comata* and *Spok2* do interact epistatically (***Grognet et al., 2014***). To determine if *Spok1* is also dominant to *Spok3* and *Spok4*, crosses were conducted between strain T_D_ and strains of *P. anserina*. Although T_D_ shows low fertility with *P. anserina* (***Boucher et al.***, *2017*), we were successful in mating T_D_ to a number of the *P. anserina* strains of the different *Psk* spore killer types (***Figure 4–source data 1 and 2***). Often only few perithecia were produced with limited numbers of asci available to count, but despite this obstacle, the crosses clearly demonstrate that T_D_ is dominant to *Psk-S* and *Psk-2*, and is mutually resistant to *Psk-5*. This result implies that *Spok1* provides resistance to all of the *Spok* homologs in *P. anserina* and is capable of killing in the presence of *Spok2* and *Spok3*, but not *Spok4*. The mutual resistance with *Psk-5* also demonstrates that *Spok4* provides resistance against *Spok1*. Additional crosses were also conducted with the *P. pauciseta* strain CBS237.71, which confirms no epistatic interactions between *Spok3* and *Spok4* in this strain (***Figure 4–source data 1and 2***). As both T_D_ and CBS237.71 have unique spore killing phenotypes, we assign them the labels *Psk-C1* and *Psk-P1*, respectively.

### An intron in the 5’ UTR is not required for spore killing

To investigate if the *Spok* genes are expressed during spore killing, we conducted an additional nine backcrosses of the S_5_ strains to S, in order to generate S_14_ backcrossed strains (see **methods**). We produced RNAseq data of self-killing S_14_ cultures and mapped the reads to the final assemblies of the parental strains. The expression of the *Spok* genes is evident in this data and supports the presence of an intron in the 5’ UTR of the *Spok* homologs (***Figure 1*** and ***Figure 1–Figure Supplement 2***). Given its conservation across the *Spok* homologs and since the *wtf* spore killer system in *S. pombe* was described to involve two alternate transcripts of the same gene (***Hu et al., 2017*; *Nuckolls et al., 2017***), the role of this intron in the *Spok3* spore killing activity was investigated. The intron was deleted in the plasmid bearing the *Spok3::PaPKS1* deletion cassette by site directed mutagenesis and the modified plasmid was used to transform the ΔKu70 Δ*Spok2* strain. Three transformants bearing the *Spok3* lacking the intron sequence (*Spok3* Δ*i*) were crossed to a Δ*Spok2* strain. As in the control cross with wild type (*wt*) *Spok3*, in which close to 100% killing was found, we observed that 109/109 of the asci contained two unpigmented spores (***Figure 4–Figure Supplement 2L***). Thus, *Spok3* Δ*i* displays wt killing activity. We conclude from this experiment that the unspliced form of *Spok3* is not required for normal killing activity, nor does the killing and resistance function via an alternatively spliced form of this intron.

### Functional annotation of SPOK3 predicts three ordered domains

In order to gain insights on the molecular function of the SPOK proteins, domain identification was performed with HHPred and a HMM profile based on an alignment of 282 *Spok3* homologs from various Ascomycota species. The SPOK3 protein was predicted to be composed of three folded domains (located at positions ∼40 – 170, 210 – 400 and 490 – 700 in the protein) separated by two unstructured domains (∼170 – 210 and 400 – 490) as shown in ***Figure 5***. No functional identification was recovered for domain 1, however a coiled-coil motif was found in the N-terminal 40 amino acids and predicted to form a parallel dimer, which corresponds to the variable length repeat of the nucleotide sequences (***Figure 1*A**). Domain 2 showed homology to a class of phosphodiesterase of the PD-(D/E)XK superfamily (∼214 – 325) with the catalytic residues forming the PD-(D/E)XK motif spanning positions 219 to 240 in the SPOK3 sequence (***Steczkiewicz et al., 2012***). The best hit in HHPred was to the HsdR subunit of a type-I restriction enzyme from *Vibrio vulnificus* (***Uyen et al., 2009***). The sequences align in the catalytic core region in the PD-(D/E)XK motif and also around a QxxxY motif (294 – 298 in SPOK3) that was found to be important for nucleic acid binding and nuclease activity (***Sisáková et al., 2008***) (***Figure 5–Figure Supplement 2***).

**Figure 5.**
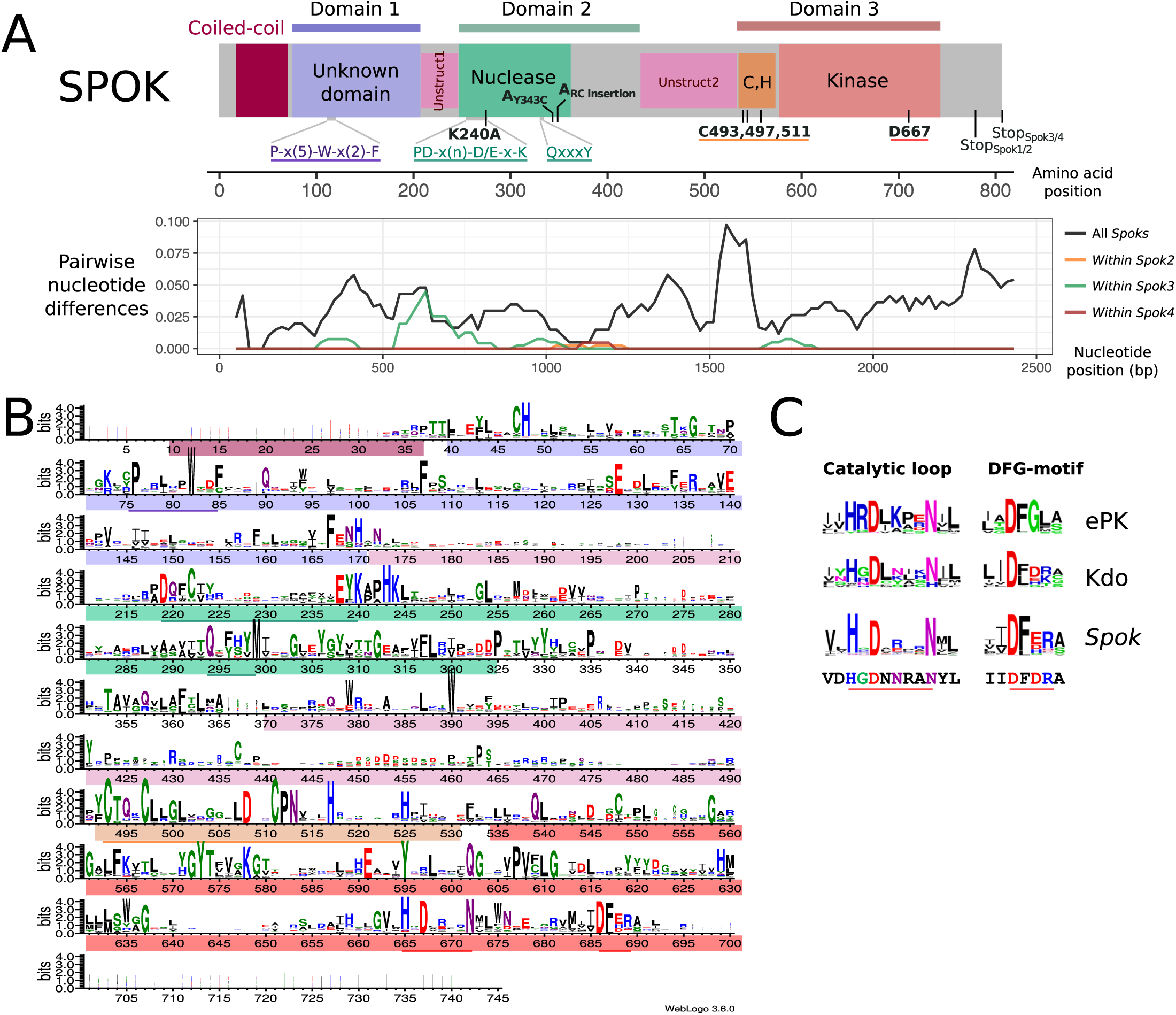
Functional annotation of the SPOK3 protein. **A** A schematic representation of a SPOK protein. Domain diagram of the SPOK3 protein displaying the N-terminal coiled-coil region (in purple), the N-terminal domain of unknown function (in dark purple), the two unstructured regions (in blue), the PD-(D/E)XK nuclease domain in green, the cysteine cluster region (in orange) and the kinase domain in red. Position of key residues and conserved motifs are given with the same color code. An amino acid length ruler in given above the diagram. A plot of the pairwise nucleotide distances among all alleles of a given *Spok* indicates which regions of the protein are conserved or divergent, and where are located the polymorphisms within a single *Spok* gene. The predicted unstructured regions generally show higher divergence. **B** HMM profile derived from an alignment of 282 SPOK3 homologs from Ascomycota showing conserved residues. The domains identified in **A** are shown with the same color code and key motifs and residues underlined. The profile was generated with Web logo v3. **C** Comparison of the HHM profiles in the catalytic loop and DFG-motif region in eukaryotic protein kinases and Kdo kinase (an ELK) (***Kannan et al., 2007***) with the same region in *Spok*-homologs. The sequence below corresponds to the SPOK3 sequence. **Figure 5–Figure supplement 1.** Visualization of an amino acid alignment for the SPOK proteins. **Figure 5–Figure supplement 2.** Model of sPOK3 domain 3. **Figure 5–source data 1.** Amino acid alignment of the SPOK proteins in the *Podospora* complex. **Figure 5–source data 2.** Gremlin amino acid alignment of SPOK proteins closely related to those in *Podospora.* **Figure 5–source data 3.** Transformation efficiency of *Spok3* manipulations.

Domain 3 was identified as a kinase domain (∼539 – 700) as predicted previously by ***Grognet et al.*** (***2014***). Additionally, a motif with a cluster of three highly conserved cysteine residues and histidine (C-x3-C-x13-C-x5-H-x7-H) reminiscent of zinc finger motifs was identified upstream of the kinase motif (***Figure 5***). As previously reported for *Spok2*, D667 was identified as the catalytic base residue in the catalytic loop (subdomain VIb) of the kinase domain. While kinases often use other proteins as substrates, they may also target small molecules (***Smith and King, 1995***). Inspection of the VIb and VII functional regions, which are informative regarding kinase substrate specificity, suggests that the *Spok*-kinase domain might be more closely related to eukaryotic-like kinases (ELKs) than to eukaryotic protein kinases (ePKs) raising the possibility that this kinase domain is not necessarily a protein kinase domain but could phosphorylate other substrates (***Steczkiewicz et al., 2012*; *Kannan et al., 2007***).

The SPOK proteins show a large degree of conservation among them and analyses of molecular evolution suggest that different domains of the protein evolve under different constraints. ***Table 3*** displays pairwise comparisons of the SPOK proteins. We tested whether any sites were evolving under positive selection using PAML 4.8 (***Yang, 2007***). The model of positive selection (M2) did not fit our data significantly better than its nested neutral model (M1). Furthermore, a likelihood test of model M3 (heterogenous site model) against the null model M0 (Homogeneous site model) showed no significant difference, which is likely due to the small number of sequences used in the analysis. In lieu of the site specific model, we calculated d_N_/d_S_ ratios for the three predicted domains. The average d_N_/d_S_ ratios of *Spok2, Spok3*, and *Spok4* are 2.70, 0.36, and 0.86 for domain 1, domain 2 and domain 3, respectively. This result suggests that domain 1 evolves under positive selection, domain 2 under purifying selection, and domain 3 under neutral or weakly purifying selection in *P. anserina*.

**Table 3.**
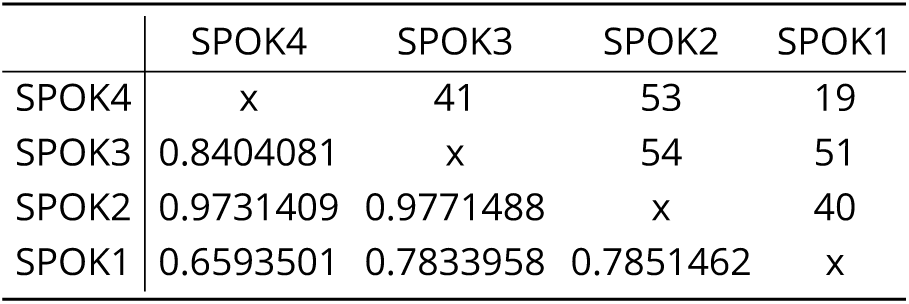
Pairwise statistics between SPOK homologs. The d_N_/d_S_ ratios, averaged across the coding region are shown below the diagonal, pairwise amino acid changes are shown above.

|  | SPOK4 | SPOK3 | SPOK2 | SPOK1 |
| --- | --- | --- | --- | --- |
| SPOK4 | x | 41 | 53 | 19 |
| SPOK3 | 0.8404081 | x | 54 | 51 |
| SPOK2 | 0.9731409 | 0.9771488 | x | 40 |
| SPOK1 | 0.6593501 | 0.7833958 | 0.7851462 | x |

### The killing and resistance functions can be attributed to separate domains

The ability of the *Spoks* to perform both killer and resistance function with a single protein is unique among meiotic drive systems (***Bravo Núñez et al., 2018***). To investigate the role that the aforementioned domains may play in these two functions, we constructed a number of point mutations and truncation variants of *Spok3* and assayed their ability to kill or provide resistance in vegetative cells. We are able to determine that domain 2 is important for killing activity while domain 3 is important for resistance activity.

It was shown previously that the kinase domain of SPOK2 (***Figure 5***) is involved in the resistance function (***Grognet et al., 2014***). We generated a point mutant affected for the predicted catalytic as-partic acid residue of *Spok3* (D667A). The mutant allele was first used in transformation of a Δ*Spok2* recipient strain. This *Spok3* D667A mutant allele leads to a drastic reduction in transformation efficiency (***Figure 4–source data 2***) while the *Spok3 wt* allele only moderately affects the number of transformants. Since this first approach results in random integration and potential multicopy insertion, we also attempted to introduce the mutant *Spok3* D667A allele as a single copy at the *PaPKS1* locus as described above for *wt Spok3*. The initial transformants were heterokaryotic and displayed sectors of abnormal growth that corresponded to unpigmented mycelium presumably containing nuclei with *Spok3* D667A that inserted at *PaPKS1*. Monokaryotic transformants could be recovered and were tested in killing activity in a cross to a Δ*Spok2*. Four-spored asci with two white and two black spores were observed, suggesting that the D667A mutation abolishes spore killing. However, when the integrated *Spok3* allele was amplified by PCR and sequenced, it appeared that the allele presents a GAG to TAG mutation leading to a premature stop codon in position 282 (E282stop). This result is consistent with the observation that *Spok3* D667A affects transformation efficiency and is toxic. Moreover, we detected expression of *Spok2* and *Spok1* in monokaryotic cultures (strains Wa63-and T_D_), suggesting that *Spok* activity is not restricted to the sexual cycle (***Figure 1–Figure Supplement 2***). No further attempts to insert the mutant allele at *PaPKS1* were made.

If toxicity of the *Spok3* D667A allele in vegetative cells is mechanistically related to spore killing, it is expected that this toxicity should be suppressed by *wt Spok3*. Therefore, we assessed whether *Spok3* D667A toxicity in vegetative cells is suppressed by co-expression with *wt Spok3*. Co-transformation experiments were set up with *Spok3* D667A used as the transformation vector in the presence or absence of *wt Spok3*. As in the previous experiment, *Spok3* D667A alone was found to affect transformation efficiency, but this effect was suppressed in co-transformations with *Spok3* (***Figure 4–source data 2***). This experiment confirms that *Spok3* D667A is only toxic in the absence of *Spok3*. Therefore, the *Spok*-related killing and resistance activities can be recapitulated in vegetative cells.

We also analyzed the role of the conserved cysteine cluster just upstream of the kinase domain. Three strains with point mutations in that region were constructed (a C493A C497A double mutant and C511A and C511S point mutants) and the mutant alleles were used in transformation assays as previously described for *Spok3* D667A. All three mutants reduced transformation efficiencies as compared to the controls and this effect was suppressed in co-transformations with *wt Spok3* (***Figure 4–source data 2***). These results suggest that the kinase domain and the cysteine-cluster region are both required for *Spok*-related resistance function but not for the killing activity. To test this, we constructed a truncated allele of *Spok3* which lacks these two regions: *Spok3*(1–490) (see ***Figure 5–Figure Supplement 1***). The *Spok3*(1–490) allele drastically reduced transformation efficiencies and this effect was suppressed in co-transformations with *wt Spok3* (***Figure 4–source data 2***). If, as proposed here, the toxicity and suppression activities assayed in vegetative cells are mechanistically related to spore killing, then domain 3 appears to be required for the resistance function but dispensable for the killing activity which can be carried out by the N-terminal region of the SPOK3 protein (domains 1 and 2).

Next we analyzed the role of the predicted nuclease domain (domain 2) in spore killing activity. We generated a point mutant affected for the predicted catalytic core lysine residue (K240A). Introduction of this point mutation in the *Spok3*(1–490) allele abolished its killing activity in transformation assays (***Figure 4–source data 2***) suggesting that the nuclease domain is required for killing activity. The *Spok3* K240A mutant was then inserted at the *PaPKS1* locus and the resulting knock-in strain was crossed with a Δ*Spok2* strain (to assay killing) and to a *Spok3::PaPKS1d* strain (to assay resistance) (***Figure 4–Figure Supplement 2M and N***). In the cross to Δ*Spok2*, no killing was observed: the majority of the asci were four-spored with two white and two black spores (308/379, 81.2%) indicating that the K240A mutation abolishes spore-killing activity of *Spok3*. In the *Spok3 K240D::PaPKS1* x *Spok3::PaPKS1d* cross, no killing was observed: the majority of the asci were four-spored with two white and two black spores (268/308, 87%). These crosses indicate that the *Spok3* K240A allele has lost killing ability but it has retained resistance. ***Grognet et al.*** (***2014***) reported that strain A bears a mutant allele of *Spok2* affected for killing but retaining resistance. The mutations in that allele fall in a conserved region of the nuclease domain (***Figure 5***) and map on predicted structural models in close vicinity of the catalytic lysine residue (K240 in SPOK3) and the other catalytic residues (***Figure 5–Figure Supplement 2***). Properties of the *Spok2* allele of strain A provide independent evidence that the nuclease domain of SPOK proteins is involved in killing activity but dispensable for resistance.

### Phylogenetic distribution of *Spok* genes

A search for closely related homologs of the *Spoks* across fungi reveals no closely related proteins among other members of the Sordariales. However, numerous species in the Hypocreales possess homologs, many of which have more than one putative copy per genome (***Figure 6***). Proteins with high similarity can also be found across other orders of the Sordariomycetes, namely the Xylariales and Glomerellales, as well as in one species of the Eurotiomycetes, *Polytolypa hystricis* (Onygenales). A maximum likelihood analysis of these sequences produced a phylogeny that can be robustly divided into two clades, one of which contains the NECHA_82228 sequence from *Nectria haematococca* (Clade I), and the other which contains the *Podospora Spok* homologs (Clade II) (***Figure 6***). NECHA_82228 was previously introduced into *P. anserina*, and the genetically modified strain produced empty asci when mated to a naïve strain, suggesting that it has a killing action (***Grognet et al., 2014***). Note that the sequences in Clade I are present in single copies per strain, except for *Fusarium oxysporum f. sp. pisi*, suggesting that they are all orthologs and hence, that the rate of gene duplications are low in this group. In contrast, many of the sequences in Clade II are present in multiple copies per strain. It is particularly notable how many *Spok* homologs are present in *F. oxysporum* and the number of copies that are found in each genome. Several of the duplicate *Spok* homologs are present on the lineage specific chromosomes of *Fusarium* that are often associated with pathogenicity (***Armitage et al., 2018***). The insect pathogens *Metarhizium rileyi* and *Cordyceps fumosorea* exhibit a number of divergent copies of *Spok* homologs with three and five copies respectively. This is in stark contrast to *Pseudomassariella vexata* and *Hirsutella minnesotensis* that have multiple, though nearly identical copies. The Clade II *Spok* homologs appear to diversify within each strain/species in much the same way as the *Spok* genes do in *Podospora*, with variable lengths of the coil-coil repeat region and frameshift mutations that relocate the stop codon. A few of the sequences may also represent pseudogenes as evident by premature stop codons and/or frameshifts, although this might also be the result of unidentified introns (***Figure 6*** and ***Figure 6–source data 1***).

**Figure 6.**
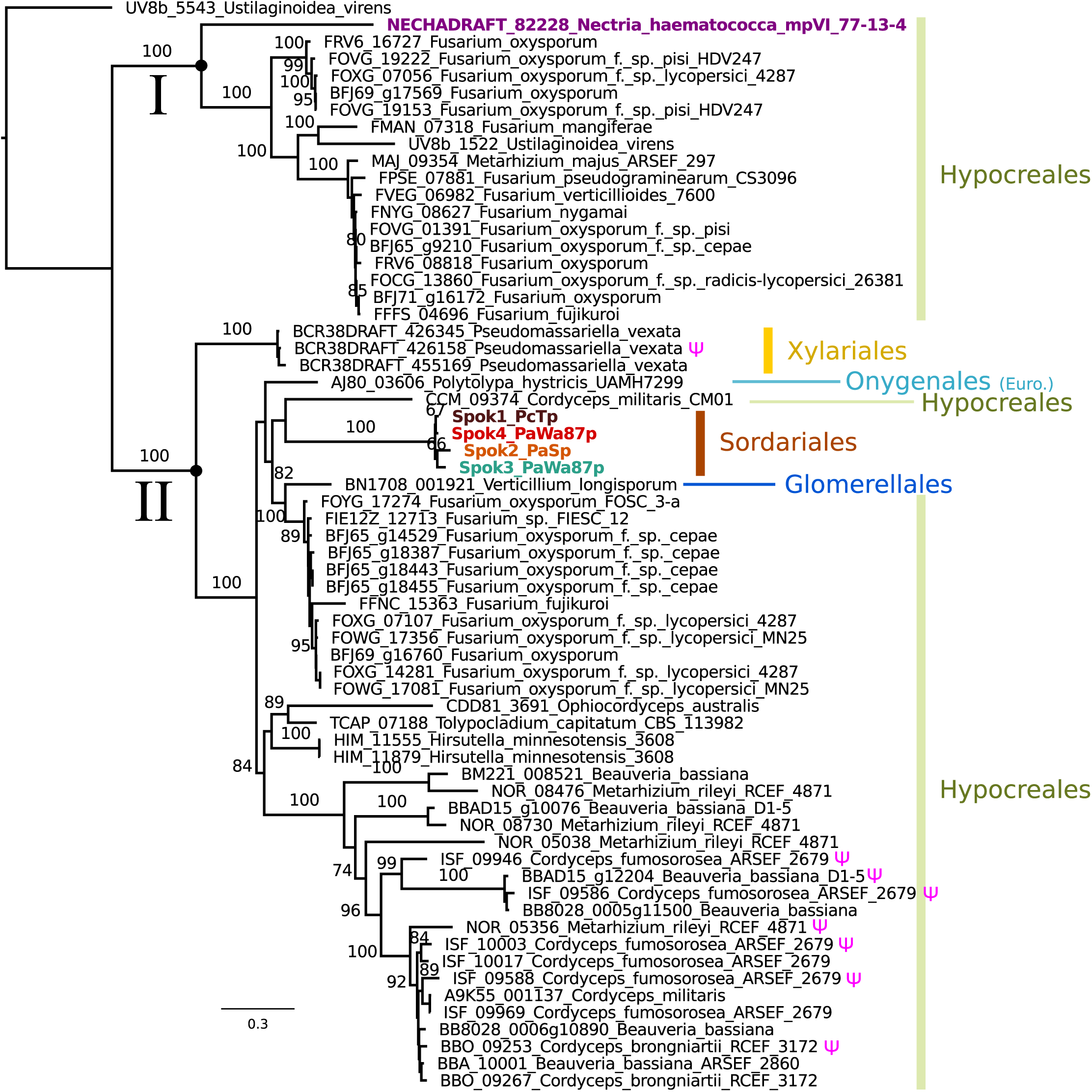
A maximum likelihood phylogenetic tree of closely related SPOK homologs. The majority of sequences come from the Hypocreales, but other lineages of Sordariomycetes are represented, as well as one species from the Onygenales (Eurotiomycetes). The clade that includes the *Podospora* SPOKs contains within-genome duplicates and has a number of putative pseudogenes (marked with a Ψ symbol). The NECHA_82228 protein (in purple) has been demonstrated to exhibit some spore killing characteristics in a *P. anserina* strain. Rooting was based on the broader alignment generated for the protein domain predictions. Bootstrap support values higher than 70 are shown above branches, which are proportional to the scale bar (substitutions per site). Tip labels follow the convention of locus name, species, and strain ID when available. **Figure 6–source data 1.** Codon-guided alignment of homologs closely related to the *Podospora* SPOKs.

## Discussion

The identification of *Spok3* and *Spok4* has allowed us to explain the genomic basis for five of the seven *Psk* spore killer types found in natural populations of *P. anserina*. By our integrative approach of genomics, molecular biology and phenotyping, we have been able to demonstrate that the multiple drive elements genetically identified in *P. anserina* are not based on different underlying molecular mechanisms and/or specific gene interactions, but rather involve combinations of closely related driver genes belonging to the same *Spok* gene family. The *Spok* genes thus appear to be responsible for all identified drive elements in *Podospora*, with the exception of the *het-s* spore killing system.

### The *Spok* Block

The presence of the complex *Spok* block presents a unique feature among the known meiotic drive systems. Often, meiotic drive elements occupy regions of suppressed recombination that span large tracts of chromosomes (***Turner and Perkins, 1979; Hammer et al., 1989; Sandler et al., 1959***) and co-occur with complex rearrangements (***Harvey et al., 2014; Silver, 1993; Dyer et al., 2007; Svedberg et al., 2018***). In these well-studied cases the elements of the drive mechanisms are encoded by separate genes within the region, and the rearrangements and suppression of recombination is expected to have evolved to ensure that the drive machinery (eg. the toxin and antitoxin genes) is inherited as one unit (***Lyttle, 1991; Bravo Núñez et al., 2018***). In *Podospora*, a single *Spok* gene is fully capable of driving, thus no region of suppressed recombination is required. Nevertheless, *Spok3* and *Spok4* are found in a large region that is not syntenic with the null allele. Hence, had the *Spok* genes not been previously identified from more placid genomic regions, the entire *Spok* block may have been misidentified as a driving haplotype with multiple interacting components. Considering that single-gene meiotic drivers might be more common than anticipated, it becomes necessary to question whether other drive systems located within complex regions and for which the genetics are not well known may also represent single gene drivers.

The relationship among the *Spoks* can provide insight as to the evolutionary history of the *Spok* block. The observation that *Spok3* and *Spok4* are both present in the *Spok* block in a duplicate region suggest that these represent homologs that formed via duplication. However, this scenario is contradicted by the finding that *Spok4* shares many features with *Spok1* of *P. comata*, yet not *Spok3*. It is possible that past hybridization between *P. anserina* and *P. comata* resulted in a transfer of *Spok4* to *P. comata* and that this gene has since diverged to become *Spok1*. In such a case, subsequent gene conversion between *Spok3* and *Spok4* would need to be invoked to explain certain features like the shared frameshift variant at the end of the CDS. If instead one assumes that the invasion of *Spok4* into *P. comata* (or of *Spok1* from *P. comata* to *P. anserina*) occurred prior to the duplication event that produced *Spok3* and *Spok4, Spok3* would have to mutate at a much higher rate than *Spok4* to explain the current pattern of divergence. Alternatively to duplication, *Spok3* and *Spok4* could be the result of divergence between different populations and ended up in their current distribution due to the fusion of two independent *Spok* blocks. Yet, another possible origin of *Spok3* and/or *Spok4* may be from another close relative, *P. pauciseta*, a scenario supported by our finding that the *P. pauciseta* strain CBS237.71 possess a *Spok* block with copies of both *Spok3* and *Spok4* that are nearly identical to the *P. anserina* alleles. Noteworthy, all possible scenarios outlined above invoke the introgression of *Spok* genes between species, most likely via hybridizations. Such interspecies interactions mediating the introgression of meiotic drive genes between species would not be a unique phenomenon to *Spok* genes of *Podospora*, as meiotic drive genes in *Drosophila* have been observed to cross species boundaries and erode barriers of reproduction (***Meiklejohn et al., 2018***). Further analyses of the genomes of populations of multiple *Podospora* species is needed in order to resolve the history of the *Spok* genes and the block.

At this stage, our data strongly suggest that the *Spok* block is moving in the genomes as a unit, but nevertheless, the mechanism of movement remains unknown. It may be hypothesized that movement of the block is achieved via an interaction with TEs at different genomic locations and non-allelic homologous recombination. This hypothesis is supported by the observation that the *Spok* genes outside of the *Spok* block, including *Spok*Ψ *1*, are not located at the same position in the different species, and that they are often surrounded by similar TEs. Such movement may be under selection as matings between strains that have the same *Spok* genes but in different locations will result in spore killing. Furthermore, due to the idiosyncrasies of meiosis in *Podospora*, the position of the block may be under selection as the killing frequency is dependent on the frequency of crossing over with the centromere. Alternatively, the TEs may simply accumulate around the *Spok* genes because of a reduced efficacy of purifying selection at regions linked to the driver genes and that their presence *per se* increases the chance of rearrangements. As such, the role that TEs play in generating complex regions associated with meiotic drive should be investigated further in order to determine their importance to the evolution of drive.

### Molecular function of the *Spoks*

Spore killing systems display analogies to toxin-antitoxin (TA) systems in bacteria and it is interesting to note that many toxin families rely on nuclease activity (***Harms et al., 2018***). The contrast between our system and TA systems, however, resides in the fact that *Spok* toxin and antitoxin activities appear to be supported by the same protein molecule. While it is premature to propose a model for the molecular basis of *Spok*-gene drive, it can be stated that the kinase activity is able to counter the toxic activity of the nuclease domain of the same protein. One may hypothesize that autophosphorylation of the SPOK proteins relieves toxicity by inhibiting the nuclease activity. Alternatively, it could be that it is the phosphorylation of a distinct macromolecule or metabolite that nullifies toxicity. This last hypothesis is supported by the fact that the kinase domain of SPOK proteins resemble small molecule kinases more than protein kinases. In a simple model, the same molecule could be the target of both the kinase and nuclease activity. One can for instance imagine that the phosphorylation of the target would make it recalcitrant to the toxic action of the nuclease domain. All killing models have to explain why the proposed inhibitory activity of the kinase domain occurs only in spores bearing the *Spok* gene, yet suicidal point mutations can be rescued in trans (***Grognet et al., 2014***). The kinase and nuclease activity of the SPOK proteins might be differentially concentration-dependent, with the kinase activity favored at high SPOK-protein concentrations presumably occurring only in spores expressing the *Spok* gene. Alternatively, the possibility for kinase activity to protect against toxic activity of the nuclease domain might be temporally constrained during ascospore maturation so that spores exposed to SPOK proteins later in development (those not bearing *Spok* genes) might not benefit from the protective action. In addition to the yet unresolved mechanistic basis of killing and resistance, the characterization of the *Spok* gene function described here poses another puzzle. Since all SPOK products have an active kinase, it is not yet known what changes in sequence confer the hierarchical interactions among some *Spok* genes or why not all SPOKs are able to provide resistance to one another. One possibility is that the cellular targets for the nuclease and kinase activity differ for the different SPOK proteins.

The coil-coiled domain is likely involved in protein-protein interactions, based on studies of similar protein domains (***van Maldegem et al., 2015***). The fact that *Spok1* and *Spok4* have the same length repeat in this domain could imply that protein-protein interactions of this domain are important for resistance, as *Spok1* and *Spok4* are mutually resistant. This model would agree somewhat with the results of reporter constructs from ***Grognet et al.*** (***2014***) that showed an N-terminal mCherry tag on *Spok2* produced empty asci. As the adjacent unknown domain has signatures of positive selection, it is possible that the functional divergence observed between the SPOK proteins is due to mutations in this portion of the protein. In this model, domain 1 might be responsible for target specificity of the nuclease (and kinase) activity. The killing action itself is expected to be universal among the *Spoks* and is supported by the fact that this entire domain of *Spok3* from T_G_ is identical to *Spok4*, yet appears to retain *Spok3* functionality. The identification of the role of the nuclease domain in killing and of the kinase domain in resistance provides a first mechanistic insight into the dual role of *Spoks*. However, further dissection of the molecular action of these proteins is required to fully understand the molecular basis of *Spok* drive.

### Absence of resistance

One of the main factors that stands out in the *Podospora* system as compared to the other well studied spore killers is the lack of resistant strains. Only one strain of *P. anserina* (strain A) has ever been described as resistant (***Grognet et al., 2014***). The point mutations of *Spok3* induced in the laboratory imply that it is trivial to create a resistant strain, since only a single nucleotide change was required. Likewise, the resistant strain A *Spok2* is different from the reference allele only by two novel insertions. As such, the lack of resistance does not appear to be the result of a mechanistic constraint. Potentially, the current *Spok* gene distribution could be a relatively young phenomenon and resistance could evolve over time. Another possibility is that resistance itself is somehow costly to the organism and selected against. Additionally, it is puzzling that none of the *Spoks* in *P. anserina* show cross resistance. Intuitively, it would seem advantageous for novel *Spok* homologs to evolve new killing functions while maintaining resistance to the other *Spok* homologs. Again, the lack of cross-resistance does not solely appear to be the result of functional constraints, as *Spok1*, which is highly similar to *Spok4*, is resistant to all other *Spok* homologs. It is possible that it is more advantageous to combine multiple independent spore killers than to have a single broadly resistant gene. This option is supported by two observations presented in this study: the occurrence of the killing hierarchy and the association of *Spok3* and *Spok4*. The fact that *Spok3* and *Spok4* are present in the *Spok* block means that they are in tight linkage with each other. It may be the case that the linkage was selected for because it provided strains with the ability to drive against strains with just *Spok3* or just *Spok4*. However, this association could also be simply the result of a duplication without invoking selection. Whether the killing hierarchy we observe in *P. anserina* is due to a complex battle among the *Spok* homologs or a result of the existence of the *Spok* block will require further experimentation and mathematical modeling to resolve.

### Evolutionary dynamics of the *Spoks*

Some interesting aspects of meiotic drive in *Podospora* identified herein bears numerous parallel features to the *wtf* genes that are responsible for drive in *S. pombe*. There is no sequence similarity or conserved domains between the *Spok* and *wtf* genes, and *Podospora* and *Schizosaccharomyces* are only distantly related (∼500 million years diverged) (***Wang et al., 2009***; ***Prieto and Wedin, 2013***). Yet these systems display similar evolutionary dynamics within their respective species. Both of these systems are built of multiple members of gene families, that appear to duplicate, rapidly diverge to the point where they no longer show cross reactions (potentially with the aid of gene conversion), and then pseudogenize and become nonfunctional (***Bravo Núñez et al., 2018***; ***Hu et al., 2017***; ***Nuckolls et al., 2017***). Both systems also have close associations with TEs (***Bowen et al., 2003***). ***Hu et al.*** (***2017***) invoke LTR-mediated non-allelic homologous recombination as a possible mechanism for *wtf* gene deletion in a lab strain of *S. pombe*. While we provide evidence for the deletion of *Spok2*, it does not fit with expectation for being LTR-mediated, but as TEs are still accumulating in the region, other TE related processes may have been involved in the deletion.

The factors determining the abundance and diversity of multigene family meiotic drivers in a species are the rates of gene duplication and loss, and time since origin. In the case of the *Spok* genes, we expect a low rate of deletion as they approach fixation, due to the dikaryotic nature of *Podospora*. Specifically, when first appearing, a deletion is only expected to be present in one of the two separate nuclear genomes maintained within a dikaryon. Any selfing event should erase (i.e. drive against) the deletion, meaning that in order to become homoallelic for a deletion, the strain would have to outcross with another individual with no *Spoks* or different *Spoks* from itself. Such outcrossing could allow deletions of *Spok3* and *Spok4*, but as *Spok2* is nearly fixed in the population, any outcrosses event should also lead to the deletion being eliminated by the driving action of *Spok2*. A possible solution to the paradoxical finding that *Spok2* appears to have been lost occasionally is that the incomplete penetrance of *Spok2* may have allowed spores that were homoallelic for the deletion to survive and persist. In this sense, *Spok2* fits the *wtf* model of driver turn over well, wherein it is beginning to lose killing function after becoming fixed in the population. *Spok*Ψ*1* is missing the portion of the gene responsible for killing and the small *Spok* fragment of *P. comata* also corresponds to the resistance part of the gene. Both these observations suggest the killing domain may have been lost prior to these genes becoming fully pseudogenized and hints that they may have functioned as resistance genes.

It has been pointed out that spore killing may be a weak form of meiotic drive, since the transmission advantage is relative to the number of spores produced in a given cross, but there is no absolute increase at the population level (***Lyttle, 1991***). Hence, a spore killer requires an additional fitness advantage to reach fixation in a population (***Nauta and Hoekstra, 1993***). It is thus striking that *Spok2* is close to fixation in at least the French and Dutch populations, bringing into question the direct fitness effects of the *Spok* genes. On the other hand, the *Spok* block (and hence *Spok3* and *Spok4*) seems to be in relatively low frequency. It is possible that the rate at which the *Spok* block switches position is higher than the rate at which the *Spoks* can sweep to fixation. As such, the dynamics of *Spok* genes within the *Spok* block might differ from the *Spok2/wtf* life-cycle and may explain why spore killing is observed to be polymorphic in *P. anserina*. Additionally, *P. anserina* is capable of selfing, which may slow down the rate of fixation of the genes. Moreover, the vegetative and/or sexual expression of *Spok* genes might be deleterious in itself, and hence natural selection might be increasing or maintaining the frequency of strains without all *Spok* homologs. Overall, this complex system requires population genetic modelling to resolve the factors affecting the frequency of the *Spok* genes in populations of this fungus.

### Evolutionary history of the *Spok* gene family

Looking more broadly at *Spok* genes across fungi for which genome sequences exist, it is rather interesting that *Spok* homologs are found in closely related orders, but not in other species of the Sordariales. This finding suggest that the *Spok* genes are transferred horizontally among evolutionarily disparate groups. This hypothesis is supported by the fact that the eurotiomycete *Polytolypa hystricis* possesses a closely related homolog to the *Podospora Spoks*. However, the phylogeny presented here shows that the homologs that group with the *Podospora Spoks* do generally agree with the known relationships among Sordariomycetes (***Maharachchikumbura et al., 2015***), suggesting that the *Spok* genes could be ancestral to the Sordariomycetes, but lost in most groups. Such a scenario would imply that there are long term consequences of possessing spore killer genes, even if they are fixed in the population.

Previously, proteins from *Nectria haematococca* and *Fusarium verticillioides* were identified as close homologs of the SPOK proteins, and it was demonstrated that the Necha_82228 protein induces spore abortion in synthetic knock-ins of *P. anserina* (***Grognet et al., 2014***). Based on diversification patterns, the phylogeny presented here suggests that the *N. haematococca* and *F. verticillioides* sequences may represent orthologs that are conserved among the Hypocreales, but do not represent meiotic drive genes since only one presumably orthologous copy is typically found. In contrast, the numerous closely related *Spok* homologs in *F. oxysporum* suggest that these genes could potentially be driving in this species. However, no sexual cycle has been observed in *F. oxysporum*. Given that we demonstrate vegetative killing with *Spok3*, it is possible that the *Fusarium Spoks* operate in vegetative tissue to ensure the maintenance of the pathogenic associated chromosomes. Alternatively, as *F. oxysporum* strains have been found with both mating type alleles (***O’Donnell et al., 2004***), there may be a cryptic sexual cycle in which the *Spok* homologs are active.

## Conclusions

With this study, we have provided a robust connection between phenotype and genotype of spore killing in *P. anserina*. We showed that meiotic drive in *Podospora spp.* is governed by genes of the *Spok* family, a single locus drive system that confers both killing and resistance within a single protein, which synergize to create hierarchical dynamics by the combination of homologs at different genomic locations. The *Spok* genes are prone to duplication, diversification and movement in the genome. Furthermore, our results indicate that they likely evolved via cross-species transfer, highlighting potential risks with the release of synthetic gene drivers for biological control invading non-target species. Moreover, we present evidence that homologs of the *Spok* genes might have similar dynamics across other groups of fungi, including pathogenic strains of *Fusarium*. Taken together, the *Spok* system provides insight into how the genome can harbour numerous independent elements enacting their own agendas and affecting the evolution of multiple taxa.

## Methods

### Fungal material

The fungal strains used in this study are listed in ***Table 1*** and were obtained from the collection maintained at the Laboratory of Genetics at Wageningen University (***van der Gaag et al., 2000***) and the University of Bordeaux. Strains with the “Wa” identifier were collected from the area around Wageningen between 1991 and 2000 (***Hermanns et al., 1995***; ***van der Gaag et al., 1998, 2000***). Strains S, Y, and Z were collected in France in 1937 (***Rizet, 1952***; ***Belcour et al., 1997***). Strain S is commonly used as a wild type reference, and an annotated genome (***Espagne et al., 2008***) is publicly available at the Joint Genome Institute MycoCosm website (https://genome.jgi.doe.gov/programs/fungi/index.jsf) as “Podan2”. It remains unclear where exactly T_D_ and T_G_ were collected, given the labelling confusion.

Representative strains for the *Psk* spore killer types from the Wageningen collection were phenotyped to confirm the interactions described by ***van der Gaag et al.*** (***2000***). Strains Wa87 and Wa53 were selected as representative of the *Psk-1* type, Wa28 for *Psk-2*, Wa21 for *Psk-3*, Wa46 for *Psk-4*,Y for *Psk-5*, Wa47 for *Psk-6*, and Wa58 for *Psk-7*. Strains S and Wa63 were used as reference strains and are annotated as *Psk-S*. Strain Wa58 mated poorly in general, so strain Z was used as a mating tester for the *Psk-7* spore killer type as well. For all crossing experiments and genome sequencing, we isolated self-sterile monokaryons (i.e., haploid strains containing only one nuclear type) from spontaneously produced 5-spored asci (***Rizet and Engelmann, 1949***), identified their mating type (mat+ or mat-) by crossing them to tester strains, and annotated them with +/-signs accordingly.

### Culture and crossing conditions

All crosses were performed on Petri-dishes with Henks Perfect barrage medium (HPM). This media is a modified recipe of PASM2 agar (van Diepeningen et al. 2008), where 5gL^−1^ of dried horse dung are added prior to autoclaving. Strains were first grown on solid minimal medium, PASM0.2. For each cross, a small area of mycelia of each of two monokaryons was excised from the plates and transferred to HPM. Perithecia (fruiting bodies) form at the interface between sexually compatible mat+ and matmonokaryons. Mature perithecia with fully developed ascospores were harvested after 8 – 11 days from which the percentage of 2-spored asci were evaluated to determine the killing percentage (***Box 1***). All cultures were incubated at 27 °C under 70% humidity for a 12:12 light/dark cycle. Barrage formation was also evaluated on HPM, whereby confrontations between mycelia of two different strains will produce a visible line of dead cells if they are vegetatively incompatible, for details see (van der Gaag, Debets, and Hoekstra 2003).

### DNA and RNA extraction and sequencing

#### Culturing, extracting and sequencing genomic DNA using Illumina HiSeq

Monokaryotic strains of *P. anserina* were grown on plates of PASM0.2 covered with cellophane. The fungal material was harvested by scraping mycelium from the surface of the cellophane and placing 80 mg to 100 mg of mycelium in 1.5ml Eppendorf tubes, which were then stored at -20 °C. Whole genome DNA was extracted using the Fungal/Bacterial Microprep kit (Zymo, www.zymo.com) and sequenced at the SNP&SEQ Technology platform (SciLifeLab, Uppsala, Sweden), where paired-end libraries were prepared and sequenced with the Illumina HiSeq 2500 platform (125bp-long reads) or HiSeq X (150bp-long reads) (***Table 1***).

#### Culturing, extracting and sequencing genomic DNA using PacBio RSII

In order to generate high molecular weight DNA suitable for sequencing using PacBio, eight strains were grown on PASM0.2 for 5 – 7 days (***Table 1***). The agar with mycelium was cut into small pieces and used as inoculum for flasks containing 200 mL 3% malt extract solution, which were then incubated on a shaker for 10 – 14 days at 27 °C. The mycelia was filtered from the flasks, cut into small pieces and ∼1g was allotted into 2ml tubes with screw-on caps, after which the tubes were stored at -20 °C. High molecular weight DNA was then extracted following the procedure described in ***Sun et al.*** (***2017***). In brief, the mycelium was freeze-dried and then macerated, and DNA was extracted using Genomic Tip G-500 columns (Qiagen) and cleaned using the PowerClean DNA Clean-Up kit (MoBio Labs). The cleaned DNA was sequenced at the Uppsala Genome Center (SciLifeLab, Uppsala, Sweden) using the PacBio RSII platform (Pacific Biosciences). For each sample, 10 kb libraries were prepared and sequenced using four SMRT cells and the C4 chemistry with P6 polymerase.

#### MinION Oxford Nanopore sequencing

DNA extraction was performed as for the PacBio sequencing, except that the mycelia was dissected to remove the original agar inocula and the DNA was purified using magnetic beads (SpeedBeads, GE) then sequenced without further size-selection. Monokaryotic samples T_G_+ and CBS237.71-were sequenced first in a barcoded run on a R9.5.1 flowcell using the Oxford Nanopore Technologies (ONT) rapid barcoding kit (1.5 μl RBK004 enzyme to 8.5 μl DNA per reaction). Due to low tagmentation efficiency, we did additional sequencing for T_G_+ using the ligation sequencing kit (LSK108, R9.4.1 flowcell). 500 ng DNA (25 ul) were mixed with 1.5ul NEB Ultra-II EP enzyme and 3.5ul NEB Ultra-II EP buffer and incubated for 10 minutes at 20 °C and 10 minutes at 65 °C before addition of 20 ul AMX adaptor, 1ul ligation enhancer, and 40 ul NEB Ultra-II ligase. After ligation the standard ONT washing and library loading protocol was followed and the sample was sequenced on a R9.4.1 flowcell. After sufficient sequencing depth had been achieved for sample T_G_, the flowcell was washed and the remaining barcoded samples were loaded to improve coverage also for sample CBS237.71. The sample Y+ yield less DNA (150 ng in 15 ul) and hence half the normal volume of adaptor was used (10 ul) and ligated using 20 ul Blunt/TA ligase for 15 minutes. Otherwise the standard protocol was followed, with sequencing done in a R9.4.1 flowcell. Basecalling and barcode split was done using Guppy 1.6 and Porechop (ONT) for all samples.

#### RNA sequencing

We generated transcriptomic data from dikaryotic strains that undergo spore killing during selfing. The S_14_ backcrosses (see below) were mated to the strain S in order to obtain killer heteroallelic spores (from 4-spore asci) that were dissected from ripe fruiting bodies (see ***Figure 1–Figure Supplement 2***). The spores were germinated in plates of PASM2 with 5gL^−1^ ammonium acetate added. Two days after germination, the culture was stored in PASM0.2 media at 4 °C to arrest growth. From that stock, we inoculated HPM plates with either a polycarbonate Track Etched 76 mm 0.1 μm membrane disk (Poretics, GVS Life Sciences, USA)(Psk1xS_5_ and Psk7xS_5_) or a cellophane layer (Psk2x_5_ and Psk5x_5_) on top. The mycelium was grown for ∼11 days and harvested for RNA extraction when the first spores were shot into the plate lid, ensuring several stages of fruiting body development. Note that *P. anserina* starts to degrade cellophane after ∼6 days, and therefore the polycarbonate membrane allows for longer growing periods. Spore killing was independently confirmed on HPM plates inoculated without a membrane. Additionally, in order to improve gene annotation, we grew the strains Wa63-and T_D_+ on a cellophane layer on HPM for 11 and7 days, respectively, to capture transcripts occurring during the monokaryotic phase.

The harvested mycelium was immediately frozen in liquid nitrogen and stored at -80 °C until RNA extraction. Next, 150 mg of frozen tissue were ground under liquid nitrogen and total RNA was extracted using RNeasy Plant Mini Kit (Qiagen, Hilden, Germany). The quality of RNA was checked on the Agilent 2100 Bioanalyzer (Agilent Technologies, USA). All RNA samples were treated with DNaseI (Thermo Scientific). Sequencing libraries were prepared using NEBNext Ultra Directional RNA Library Prep Kit for Illumina (New England Biolabs). The mRNA was selected by purifying polyA+ transcripts (NEBNext Poly(A) mRNA Magnetic Isolation Module, New England Biolabs). Finally, paired-end libraries were sequenced with Illumina HiSeq 2500 at the SNP&SEQ Technology platform.

### Reads processing and genome assembly

For both DNA and RNA Illumina HiSeq reads, adapters were identified with cutadapt v. 1.13 (***Martin, 2011***) and then trimmed using Trimmomatic 0.36 (***Bolger et al., 2014***) with the options `ILLUMINACLIP:adapters.fasta:1:30:9 LEADING:20 TRAILING:20 SLIDINGWINDOW:4:20 MINLEN:30`. Only filtered reads with both forward and reverse were kept for downstream analyses. For short-read mapping, we used BWA v. 0.7.17 (***Li and Durbin, 2010***) with PCR duplicate marking of Picard v. 2.18.11 (http://broadinstitute.github.io/picard/), followed by local indel re-aligning implemented in the Genome Analysis Toolkit (GATK) v. 3.7 (***Van der Auwera et al., 2013***). Mean depth of coverage was calculated with QualiMap v.2.2 (***Okonechnikov et al., 2016***).

The raw PacBio reads were filtered and assembled with the SMRT Analysis package and the HGAP 3.0 assembler (***Chin et al., 2013***). The resulting assembly was error-corrected (polished) with Pilon v. 1.17 (***Walker et al., 2014***) using the mapped filtered Illumina reads of the same strain. The samples sequenced with MinION were assembled using Minimap2 v. 2.11 (***Li, 2018***) and Miniasm v. 0.2 (***Li, 2016***), polished twice with Racon v. 1.3.1 (***Vaser et al., 2017***) using theMinION reads, and further polished for five consecutive rounds of Pilon v. 1.22 using the Illumina reads as above. Additionally, DNA Illumina reads were assembled de novo for each sample using SPAdes v. 3.12.0 (***Bankevich et al., 2012***; ***Antipov et al., 2015***) using the k-mers 21,33,55,77 and the `–careful` option. Assemblies were evaluated using QUAST v. 4.6.3 (***Mikheenko et al., 2016***). Scaffolds were assigned chromosome numbers based on homology with Podan2. BLAST searches of the scaffolds in the final assembly of the strain CBS237.71 revealed contamination by a *Methylobacterium sp.* in the MinION data (but not in the Illumina data set). The scaffolds matching the bacterium were removed from the analysis.

The assembly of the *Spok* block was visually inspected by mapping the long reads (using Minimap2) and the short reads (BWA) as above into the long-read polished assemblies. Since the MinION assemblies maintain some degree of sequencing error at repetitive regions that cannot be confidentially polished, we also assembled both types of reads into a hybrid assembly using SPAdes (same options as above) and, whenever different on short indels or SNPs but fully assembled, the sequence of the *Spok* genes was taken from the (low-error) hybrid assembly. The assembly of the *Spok* block of the T_G_+ strain was particularly challenging since the recovered MinION reads were relatively short. However, a few (<10) reads were long enough to cover the tandem duplication that contains *Spok3* (albeit with high nucleotide error rate in the assembly). The hybrid SPAdes assembly collapsed the duplication into a single copy. We therefore mapped the short reads into the hybrid assembly, confirming that the *Spok3* gene had doubled coverage and no SNPs, as expected from a perfect duplication.

Alignments of the assembled genomes were performed with the NUCmer script of the MUMmer package v. 4.0.0beta2 (***Kurtz et al., 2004***) using options `-b 200 -c 2000 -p –maxmatch`. The figures showing alignments of the *Spok* block and the *Spok2* region (***Figure 2, Figure 3***, and ***Figure 2–Figure Supplement 1***) were generated by extracting the regions from each de novo assembly and aligning them in a pairwise fashion using MUMmer as described above. The MUMmer output were then visualized using a custom Python script.

### Genome annotation

For annotation, we opted for gene prediction trained specifically on *P. anserina* genome features. We used the ab initio gene prediction programs GeneMark-ES v. 4.32 (***Lomsadze et al., 2005***; ***Ter-Hovhannisyan et al., 2008***) and SNAP release 2013-06-16 (***Korf, 2004***). All the training process was done on the sample Wa28-, for which all chromosomes were assembled (see **Results**). The program GeneMark-ES was self-trained with the script `gmes_petap.pl` and the options `–fungal –max_intron 3000 –min_gene_prediction 120`. SNAP was trained as instructed in the tutorial of the MAKER pipeline v. 2.31.8 (***Holt and Yandell, 2011***), in (***Campbell et al., 2014***), and in the SNAP README file. First, we use the Podan2 transcripts and protein models as sole evidence to infer genes with MAKER (option est2genome=1) and then we had a first round of SNAP training. The resulting HMM file was used to re-run MAKER (est2genome=0) and to re-train SNAP, obtaining the final HMM training files.

A library of repetitive elements was constructed by collecting the reference *P. anserina* transposable elements described in ***Espagne et al.*** (***2008***) available in Genbank, and combining them with the fungal portion of Repbase version 20170127 (***Bao et al., 2015***), as well as the *Neurospora* library of ***Gioti et al.*** (***2013***). In order to produce transcript models we used STAR v. 2.6.1b (***Dobin et al., 2013***) with maximum intron length of 1000 to map the RNAseq reads of all samples, followed by processing with Cufflinks v. 2.2.1 (***Trapnell et al., 2010***). For the final genome annotation, we used MAKER v. 3.01.02 along with GeneMark-ES v. 4.33, SNAP release 2013-11-29, RepeatMasker v. 4.0.7 (http://www.repeatmasker.org/), BLAST suit 2.6.0+ (***Camacho et al., 2009***), Exonerate v. 2.2.0 (***Slater and Birney, 2005***), and tRNAscan-SE v. 1.3.1 (***Lowe and Eddy, 1997***). After preliminary testing, we chose the transcripts of Psk7xS_14_ (mapped to the PacBio assembly of Wa58-) and Wa63-(PacBio assembly of the same strain) as EST evidence, and the Podan2 and T_D_ (***Silar et al., 2018***) models as protein evidence. The MAKER models of relevant regions were manually curated by comparing with RNAseq mapping and coding sequences (CDS) produced with TransDecoder v. 5.5.0 (***Haas et al., 2013***) on the Cufflinks models.

We used blastn to localize possible copies of *Spok* genes in all genome assemblies. The *Spok2* (Pa_5_10) gene from ***Grognet et al.*** (***2014***) was selected as query. We named the new *Spok* genes (*Spok3* and *Spok4*) arbitrarily based on sequence similarity, as reflected in the Phylogenetic analyses (see below). Note that the existence of *Spok3* had previously been hypothesised by ***Grognet et al.*** (***2014***), however no DNA sequence was provided. Moreover, the strain Y, in which they identified it, contains both *Spok3* and *Spok4*.

### Introgressions of the Spore-killing loci

Backcrossed strains of the various spore killer phenotypes were generated through five recurrent backcrosses to the reference strain S (S_5_) by ***van der Gaag et al.*** (***2000***). In the original study, the strains selected as spore killer parents were Wa53+ for *Psk-1*, Wa28-for *Psk-2*, Y+ for *Psk-5*, and Wa58-for *Psk-7*. The S_5_ strains are annotated as Wa170 (*Psk-1*), Wa130 (*Psk-2*), Wa200 (*Psk-5*), and Wa180 (*Psk-7*) in the Wageningen Collection, however for the sake of clarity we refer to them as Psk1xS_5_, Psk2xS_5_, Psk5xS_5_, and Psk7xS_5_.

We sequenced the S_5_ strains along with the reported parental strains using Illumina HiSeq 2500. We mapped the reads to Podan2 as described above, followed by SNP calling using the HaplotypeCaller pipeline of GATK (options: `-ploidy 1 -newQual -stand_call_conf 20.0`). We removed sites that had missing data, that overlapped with repeated elements as defined by RepeatMasker, or where all samples were different from the reference genome, using VCFtools v. 0.1.16 (***Danecek et al., 2011***), BEDtools v. 2.27.1 (***Quinlan and Hall, 2010***), and BCFtools v. 1.9 (***Danecek and McCarthy, 2017***), respectively. We plotted the density of filtered SNPs across the genome with the R packages vcfR (***Knaus and Grünwald, 2016***) and poppr (***Knaus and Grünwald, 2016***; ***Kamvar et al., 2015***). A full Snakemake (***Köster and Rahmann, 2018***) pipeline can be found at https://github.com/johannessonlab/SpokPaper. Notice that we sequenced both monokaryons of our strain S to account for the mutations that might have had occurred since the separation of the reference S strain in the laboratory of ***Espagne et al.*** (***2008***) and our S strain from the Wageningen Collection. These mutations should be present in the backcrosses, but they are independent from the spore killer elements.

Inspection of the introgressed tracks revealed that the variants of the backcross Psk1xS_5_ do not match perfectly Wa53+ (the reported parent). Given that the *Spok* content is the same as Wa53+, the introgressed track co-occurs with the expected position of the *Spok* block in chromosome 3, and the fact that the phenotype of this backcross matches a *Psk-1* spore-killer type, we concluded that Wa170 (Psk1xS_5_) in the collection actually belongs to another of the *Psk-1* backcrosses described in the doctoral thesis of ***van der Gaag*** (***2005***), likely backcrossed from Wa52. Puzzling, an introgressed track in the chromosome 3 of the Psk2xS_5_ strain does not match the expected parent (Wa28) either, both in SNPs and *het* genes alleles (***Figure 2–Figure Supplement 3***). However, other tracks in different chromosomes, including that of chromosome 5 where the *Psk-2 Spok* block can be found, do match Wa28. Likewise, Psk2xS_5_ only has *Spok2* and *Spok3* copies like Wa28. Hence, we concluded that our results are not affected by these inconsistencies.

As reported by ***van der Gaag et al.*** (***2000***), the S_5_ strains were generated by selecting ascospores from 2-spored asci of crosses between S and the spore killer parent. This procedure ensures that the offspring will be homozygous for alleles of the spore-killer parent from the spore killing locus to the centromere (***Box 1*** and ***Figure 2–Figure Supplement 3***). To eliminate as much background as possible from the spore killer parents in the backcrossed strains, nine additional backcrosses were conducted where ascospores were selected from 4-spored and 2-spored asci in alternating generations. Ascospores from the final generation were selected from 2-spored asci to ensure the strains would be homozygous at the spore-killing locus. These strains are the result of 14 backcrosses to S (S_14_) and are annotated as Psk1xS_14_, Psk2xS_14_, Psk5xS_14_, and Psk7xS_14_. The S_5_ and S_14_ strains were phenotyped by crossing the strains to their parents as well as other reference spore killer strains to confirm that the killing phenotypes remained unchanged after the backcrosses.

### Knock-out of *Spok2*

To knock-out *Spok2*, a 459 bp and a 495 bp fragment flanking the *Spok2* ORF downstream and upstream were obtained by PCR and cloned flanking the *hph* gene in the SKhph plasmid as blunt end fragments in a *Eco*RV site and a *Sma*I site. The deletion cassette was then amplified by PCR and used to transform a ΔKu70 strain (***El-Khoury et al., 2008***). Five transformants were screened for integration of the *hph* marker at *Spok2* by PCR and crossed to s. To purify the Δ*Spok2* nuclei, a heterokaryotic binucleated Δ*Spok2*/*Spok2* spore was recovered in a 2-spored ascus and use to fertilize the initial Δ*Spok2* transformant (which may or may not be heterokaryotic). Uninucleated *hygR* resistant spores were then recovered from this cross.

### Construction of a disruption cassette to insert *Spok3* or *Spok4* in the *PaPKS1* locus

To replace the ORF of the centromere-linked Pa_2_510 (*PaPKS1*) gene by one of the *Spok3* or *Spok4* genes (see **Results**), a disruption cassette was constructed as follows. A DNA fragment corresponding to the 700 bp upstream region of the *PaPKS1* ORF was amplified with oligonucleotides 5’ tcgccgcggGCTAGGGGGTACTGATGGG 3’ and 5’ cacgcggccgcCTTGGAAGCCTGTTGACGG 3’ (capital letters correspond to *P. anserina* genomic DNA sequences) and cloned in SKpBluescript vector (Stratagene) containing the nourseothricin-resistance gene *Nat* in the *Eco*RV site (vector named P1) using the *Sac*II/*Not*I restriction enzymes (upstream from the *Nat* gene) to produce the P1UpstreamPKS1 vector. Then the 770 bp downstream of the *PaPKS1* ORF was amplified with oligonucleotides 5’ tcgaagcttACAACAGTCATACGAACATG 3’ and 5’ gcggtcgacGGTACAATACGCCCTCAGTG 3’ and cloned in the P1UpstreamPKS1 vector using the *Hind*III/*Sal*I restriction enzymes (downstream from the *Nat* gene) to produce the P1UpstreamDownstreamPKS1 vector. Finally the *Spok3* and *Spok4* genes were amplified respectively from the Wa28 strain with oligonucleotides 5’ tcggcggccgcCACAGGAGCAGAGCTACGAC 3’ and 5’ gcgtctagaATATTTGGGTACTTGGCGGC 3’ and from the Wa87 strain with oligonucleotides 5’ tcggcggccgcCACAGGAGCAGAGCTACGAC 3’ and 5’ gcgtctagaCAAGGTGCCCGTGGAGTAAG 3’ and cloned in the P1UpstreamDownstreamPKS1 vector using the *Not*I/*Xba*I restriction enzymes (between *PaPKS1* upstream region and *Nat* gene) to produce the P1UpstreamDownstreamPKS1_Spok3 or the P1UpstreamDownstreamPKS1_Spok4 vector so that the *Spok3*/*Spok4* and *Nat* genes are flanked by the upstream and downstream regions of *PaPKS1* ORF to allow *PaPKS1* ORF replacement by homologous recombination. The *Spok3* and *Spok4* amplified *Spok* genes contain the ORFs flanked with 983 bp upstream the start codon and 460 bp downstream the stop codon for *Spok3*, and 984 bp upstream the start codon and 393 bp downstream the stop codon for *Spok4*, allowing expression of *Spok* genes using their native promoter and terminator regions. The disruption cassette were then amplified from the final vectors using the most distal oligonucleotides 5’ tcgccgcggGCTAGGGGGTACTGATGGG 3’ and 5’ gcggtcgacGGTACAATACGCCCTCAGTG 3’ and named PKS1∷Spok3_nat-1 and PKS1∷Spok4_nat-1.

*P. anserina* Δ*Spok2* (ΔPa_5_10) strain was obtained after disruption of the gene Pa_5_10 and replacement of its ORF with the hygromycin-resistance gene *hph* in a ΔKu70 strain. This strain was used as recipient strain for the disruption cassettes. We used 5ul of the cassettes for transfection and Nourseothricin resistant transformants were selected. As expected, most of the transformants were unpigmented and corresponded to insertion of *Spok3* or *Spok4* by replacement of *PaPKS1*. Gene replacement was verified by PCR.

### Protein annotation methods

Prediction of unstructured regions was performed in SPOK3 with PrDOS with a 2% false positive setting (***Ishida and Kinoshita, 2007***). Coiled-coil prediction was performed with LOGICOIL (***Vincent et al., 2012***), CCHMM_PROF (***Bartoli et al., 2009***) and Multicoil2 (***Wolf et al., 1997***). Domain prediction was performed using Gremlin (***Balakrishnan et al., 2011***) and RaptorX contact predict (***Ma et al., 2015***). Conserved residues were identified using Weblogo 3 (***Crooks et al., 2004***) witha Gremlin generated alignment as input. Domain identification was done with HHPred (***Zimmermann et al., 2018***).

In order to compare the diversity at the nucleotide level with the protein models, we calculated the average pairwise nucleotide differences (***Nei and Li, 1979***) for each bi-allelic site (correcting by the number of sites (n/(n -1)) while ignoring sites with gaps) on a *Spok* alignment (see below), using overlapping windows of 100 bp and steps of 20 bp. This was performed on a selected representative of each *Spok* homolog (*Spok2* of S, *Spok3* and *Spok4* of Wa87, and *Spok1* from T_D_), or for all the alleles of each *Spok* within the *P. anserina* strains.

Values of d_N_/d_S_ were calculated using the seqinr package in R (***Charif and Lobry, 2007***). Alignments were manually trimmed to calculate separate values for each domain. Tests of sequence evolution were conducted with paml 4.8 (***Yang, 2007***) using a star phylogeny of the *Spok* sequences.

### Phylogenetic analyses

The final gene models of all the *Spok* genes in *Podospora spp.* were aligned along with the sequences of *Spok2* and *Spok1* from ***Grognet et al.*** (***2014***) using MAFFT online version 7 (***Katoh et al., 2017***) with default settings (only one copy of *Spok3* from T_G_ was used). The resulting alignment was manually corrected taking into account the reading frame of the protein. Since the UTRs seem to be conserved between paralogs, 654 (5’ end) and 250 (3’ end) bps of the flanking regions with respect to *Spok2* were also included in the alignment. An unrooted split network was constructed in SplitsTree4 v. 4.14.16, build 26 Sep 2017 (***Huson and Bryant, 2006***) with a NeighborNet (***Bryant and Moulton, 2002***) distance transformation (uncorrected distances), and an EqualAngle splits transformation. SplitsTree4 was used likewise to perform a Phi test for recombination (***Bruen et al., 2006***) using a windows size of 100 and k = 6. Additionally, we used the BlackBox of RAxML-NG v. 0.6.0 (***Kozlov et al., 2018***) to infer Maximum Likelihood phylogenetic trees of the nucleotide alignment of the 5’ UTR, the coding sequence (CDS), and the 3’ UTR of the *Spok* homologs. We ran RAxML-NG with 10 parsimony and 10 random starting trees, a GTR + GAMMA (4 categories) substitution model, and 100 bootstrap pseudo-replicates for each analysis.

In order to create a phylogeny of proteins closely related to the *Spok* genes in *Podospora* (and hence likely to be meiotic drivers), the protein sequence of *Spok1* was used as a query against the NCBI genome database. We collated all hits with e-values lower than Necha2_82228, which has been shown to have some spore killing functionality in *P. anserina* previously (***Grognet et al., 2014***), with hit coverage greater than 75%, and no missing data (Ns) in the sequence. The sequences were aligned using the codon-aware program MACSE v. 2.03 (***Ranwez et al., 2018***), with the representative *Podospora Spoks* set as “reliable” sequences (`-seq`), and the rest as “non reliable” (`-seq_lr`). Many of the original gene models predict introns in the sequences, however no divergent regions were apparent in the alignment and, even if present, MACSE tends to introduce compensatory frame shifts. As such the entire gene alignment was used for the analysis. The resulting nucleotide alignment was corrected manually, translated into amino acids, and trimmed with TrimAl v. 1.4.1 (***Capella-Gutiérrez et al., 2009***) using the `-gappyout` function. A Maximum likelihood tree was then produced using IQ-TREE v. 1.6.8 (***Kalyaanamoorthy et al., 2017***; ***Nguyen et al., 2015***) with extended model selection (`-m MFP`) and 1000 standard bootstrap pseudo-replicates. The protein sequence UV8b_5543 of *Ustilaginoidea virens* was selected as outgroup based on a BioNJ tree made with SeaView v. 4.5.4 (***Gouy et al., 2010***) of the Gremlin alignment described above.

### Pool-sequencing of *Psk-1* vs *Psk-5* progeny

In order to confirm that *Spok2* is responsible of the killing between *Psk-5* and *Psk-1*, we conducted a cross between the strains Wa87 and Y. When perithecia started shooting spores, we replaced the lid of the cross plate with a water-agar plate upside-down, and let it sit for around an hour. Since *P. anserina* spores from a single ascus are typically landing together, it is possible to distinguish spores that came from an ascus with no killing (groups of four spores) from those that had killing (groups of two). To improve germination rates, we scooped spore groups of the same ascus type and deposited them together in a single plate of germination medium. After colonies became visible, they were transferred into a PASM2 plate with a cellophane layer where they grew until DNA extraction, followed by pool-sequencing with Illumina HiSeq X. In total 21 2-spore groups, and 63 4-spore groups were recovered.

The resulting short reads were quality controlled and mapped to Podan2 as above. We used GATK to call variants from the parental strains (treated as haploid) and the two pool-sequencing databases (as diploids). We then extracted SNPs, removed sites with missing data, and attempted to quantify the coverage frequency of the parental genotypes for each variant. The expectation was that spore-killing (2-spore asci) would result in a long track of homozygosity (only one parental genotype) around *Spok2*, as compared to the fully heterozygous 4-spore asci. A full Snakemake pipeline is available at https://github.com/johannessonlab/SpokPaper.

## Author contributions

**Aaron A. Vogan** Uppsala University, Uppsala, Sweden

**Contributions:** Conceptualization, Methodology, Validation, Formal analysis, Investigation, Writing - original draft, Visualization

**Contributed equally with:** S. Lorena Ament-Velásquez

**S. Lorena Ament-Velásquez** Uppsala University, Uppsala, Sweden

**Contributions:** Conceptualization, Methodology, Software, Validation, Formal analysis, Investigation, Data curation, Writing - review and editing, Visualization, Funding acquisition

**Contributed equally with:** Aaron A. Vogan

**Alexandra Granger-Farbos** Non-Self Recognition in Fungi, Institut de Biochimie et de Génétique Cellulaire, CNRS UMR 5095, Université de Bordeaux, Bordeaux, France

**Contributions:** Investigation

**Jesper Svedberg** Uppsala University, Uppsala, Sweden

**Contributions:** Conceptualization, Investigation, Writing - review and editing, Visualization

**Eric Bastiaans** Uppsala University, Uppsala, Sweden Laboratory of Genetics, Wageningen University, Wageningen, Netherlands

**Contributions:** Conceptualization, Investigation, Writing - review and editing

**Alfons J. M. Debets** Laboratory of Genetics, Wageningen University, Wageningen, Netherlands

**Contributions:** Conceptualization, Resources, Writing - review and editing

**Virginie Coustou** Non-Self Recognition in Fungi, Institut de Biochimie et de Génétique Cellulaire, CNRS UMR 5095, Université de Bordeaux, Bordeaux, France

**Contributions:** Investigation

**Hélène Yvanne** Non-Self Recognition in Fungi, Institut de Biochimie et de Génétique Cellulaire, CNRS UMR 5095, Université de Bordeaux, Bordeaux, France

**Contributions:** Investigation

**Corinne Clavé** Non-Self Recognition in Fungi, Institut de Biochimie et de Génétique Cellulaire, CNRS UMR 5095, Université de Bordeaux, Bordeaux, France

**Contributions:** Investigation, Writing - review and editing, Supervision

**Sven J. Saupe** Non-Self Recognition in Fungi, Institut de Biochimie et de Génétique Cellulaire, CNRS UMR 5095, Université de Bordeaux, Bordeaux, France

**Contributions:** Conceptualization, Methodology, Formal analysis, Investigation, Writing - review and editing, Visualization, Supervision

**Hanna Johannesson** Uppsala University, Uppsala, Sweden

**Contributions:** Conceptualization, Writing - review and editing, Supervision, Project administration, Funding acquisition

## Acknowledgments

We would like to thank Magdalena Grudzinska-Sterno for valuable assistance with DNA and RNA extractions as well as library preparations. We acknowledge support of the National Genomics Infrastructure (NGI) / Uppsala Genome Center for assistance with massive parallel sequencing. We are also thankful to Ola Wallerman for assistance with MinION Oxford Nanopore sequencing. The computations were performed on resources provided by SNIC through Uppsala Multidisciplinary Center for Advanced Computational Science (UPPMAX) under Project SNIC 2017/1-567. This study was founded by a European Research Council grant under the program H2020, ERC-2014-CoG, project 648143 (SpoKiGen), funding from The Swedish Research Council (VR) (to HJ), and by the Lars Hierta Memorial Foundation and The Nilsson-Ehle Endowments of the Royal Physiographic Society of Lund (to SLAV).

## Data availability

The full CDS sequence and UTRs of *Spok3, Spok4*, and *Spok*Ψ*1* (strain Wa87+) were deposited in NCBI GenBank under the accession numbers MK521588, MK521589, and MK521590, respectively. Raw sequencing reads were deposited on the NCBI SRA archive under the BioProject PRJNAXXXXXX.

### Appendix 1

#### The biology of *Podospora*

The life cycle of *P. anserina* is an important factor to consider when discussing the meiotic drive of the *Sp ok* genes. Although it has haploid nuclei, *P. anserina* maintains a dikaryotic (n+n) state throughout its entire lifecycle. Haploid nuclei of different mating-type are shown as white and black circles within fungal cells. The fruiting body (perithecium) is generated from dikaryotic (n+n) mycelia, usually from a single individual strain. Within the perithecium, the sexual cycle is completed to produce four dikaryotic ascospores per ascus. Occasionally, atypical spore formation may occur and result in the production of five spores in an ascus, of which two are small and monokaryotic (n). These are self-sterile and need to outcross either with a monokaryotic individual of the opposite mating type or with a dikaryotic individual to complete the life cycle. Note that outcrossing may occur via mating between either siblings or unrelated individuals of the opposite mating type. The monokaryotic spores are useful for generating self-sterile (haploid) cultures for the purposes of sequencing and laboratory mating. This intricate lifecycle is maintained a strict meiotic process.

**Appendix 1 Figure 1.**
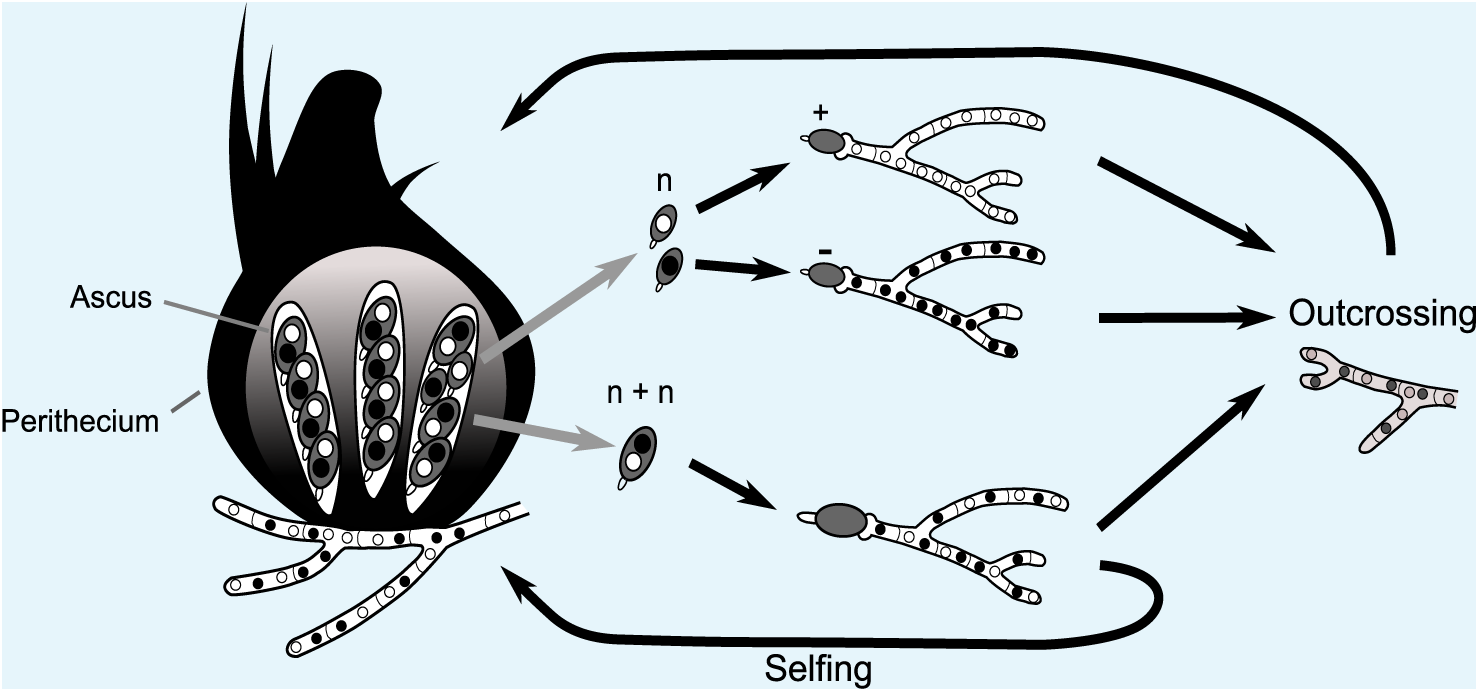
Simplified life cycle of *P. anserina*.

#### Two-locus spore killing interaction

The interaction between *Psk-1* and *Psk-7* provides a good example of how the meiotic drive dynamics of *P. anserina* result in killing even though both *Psk-1* and *Psk-7* possess the same *Spok* homologs. The three *Spok* homologs (*Spok2, Spok3*, and *Spok4*) are all present in both *Psk-1* and *Psk-7*. The observed mutual resistance is thus due to the fact that the *Spok* block (with *Spok3* and *Spok4*) is located on different chromosomes. Because chromosomes segregate independently at meiosis the expected killing percentage can be calculated as:

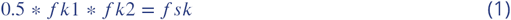

where 0.5 is due to independent assortment of chromosomes, *f k*1 is the killing percentage of strain 1, *f k*2 is the killing percentage of strain 2, and *f sk* is the spore killing frequency observed between the two strains. For *Psk-1* crossed to *Psk-7* this equals 0.27. This agrees well with the observed killing percentage of 23 – 27% (***Figure 4–Figure Supplement 1***).

### Appendix 2

#### History of *Spore* killer research in *Podospora*

Throughout the history of *spore* killing research in *Podospora*, a number of observations have been made along with corresponding hypotheses. The discovery of *Spok3* and *Spok4* provides us with the opportunity to reinterpret these data in light of the results presented herein. Here we will address data from four important works: ((***Padieu and Bernet, 1967***; ***van der Gaag et al., 2000, 2003***; ***Hamann and Osiewacz, 2004***).

#### Inconsistencies among the *Psk* designations

Our phenotyping is in accordance with the results of (***van der Gaag et al., 2000***) for strains Wa28, Wa53, Wa58, Wa63, Wa87, S, and Z, while contradictions were observed for Wa21, Wa46, Wa47, and Y. Strain Wa21 was previously categorized as *Psk-3* which is typified by inconsistent spore killing with *Psk-S* strains. Here we observed stable percentages and thus consider Wa21 to be representative of *Psk-2*. The role of *Psk-3* as a spore killer has been in doubt since its description (***van der Gaag et al., 2000***). This is in part due to the fact that ascospores are not fully aborted as for the other spore killer types. Instead small transparent ascospores can still be observed within the ascus. Here we were unable to find support for this spore killer type and it has no clear correlation between its phenotype and any *Spok* genes. We therefore find it likely that the effect is due to other incompatibility factors rather than meiotic drive.

We did not observe any spore killing in crosses between Wa46 (*Psk-4*) and Wa47 (*Psk-6*) as reported in van der Gaag 2000. Two other strains had been annotated as *Psk-6*, Wa89 and Wa90, but no other strains were recorded as *Psk-4*. Unfortunately we were not able to phenotype these strains and so we are unable to evaluate *Psk-6* further in this study. In addition, results from crosses of *Psk-4* with a *Psk-S* strain (Wa63) reveals that there is a dominance interaction between them with *Psk-S* killing *Psk-4*, which is the opposite of what was proposed in ***van der Gaag et al.*** (***2000***), i.e. that *Psk-S* kills *Psk-4*. Potentially, the original interpretation was hindered by poor mating of the *Psk-4* strain with tester *Psk-S* strains. Previously, strain Y was reported to have mutual resistance with *Psk-1*, be susceptible to *Psk-7*, and dominant over all other types. Here we report that Y is susceptible to *Psk-1* and *Psk-7*, and has mutual killing with all other types, except for crosses with naïve strains where it is dominant.

#### Allorecognition (***het*) genes and spore killing**

As the *het-s* gene is capable of causing both vegetative incompatibility and spore killing, it was hypothesized that the *Psk* loci may be as well. The S_5_ strains all demonstrate barrage formation (symptomatic of vegetative incompatibility) with strain S (***van der Gaag et al., 2003***). However when additional backcrosses were performed to generate S_14_ strains, no barrages were observed (***Figure 1***). This indicates that the spore killing types do not directly affect vegetative incompatibility or vice versa, but may be linked to loci which do. Note that the S_5_ strains contain multiple genomic regions that are not isogenic with S, some of which contain known allorecognition genes (***Figure 2–Figure Supplement 3***).

**Appendix 2 Figure 1.**
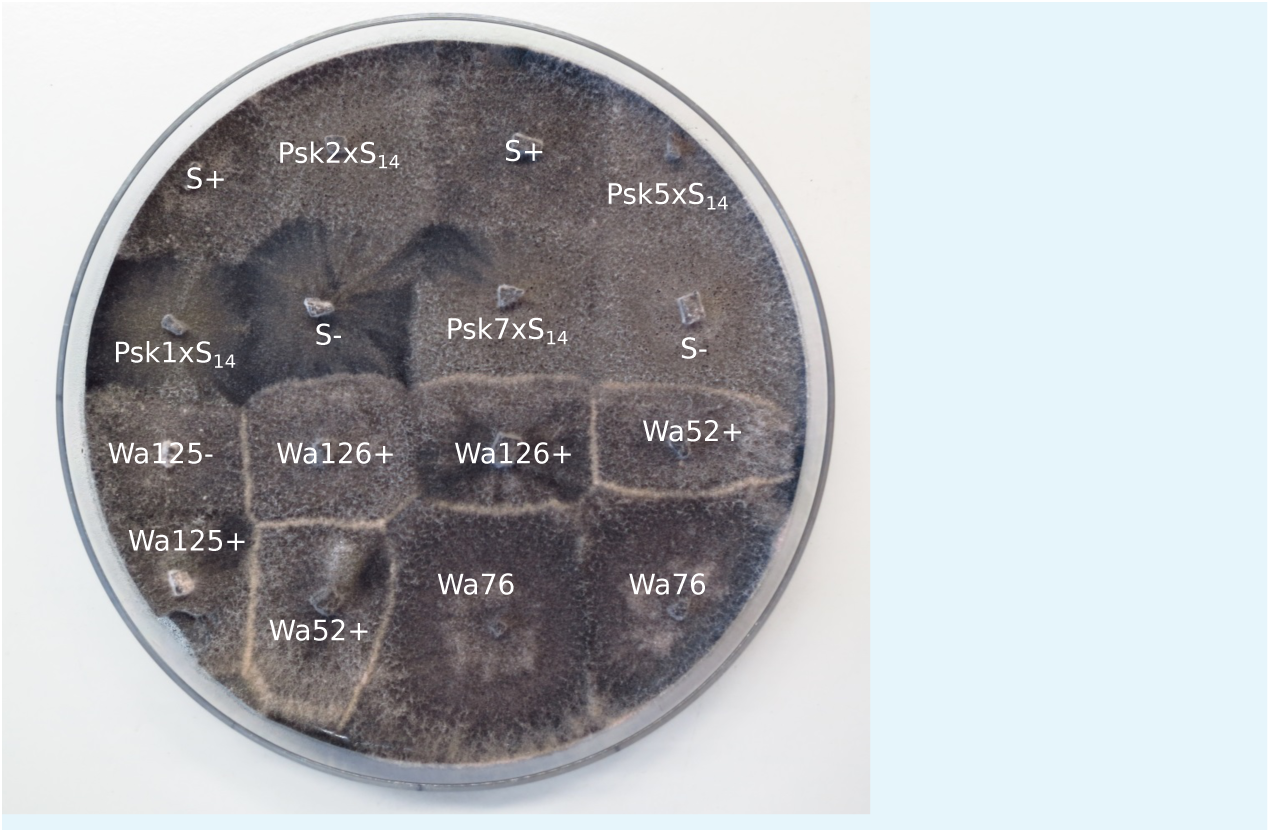
Barrage tests of the S_14_ strains. Strains Wa126, Wa76, Wa52, and Wa125 are wild isolates of *P. anserina* in the Wageningen collection. The thick white lines of mycelia demonstrate a barrage, which is indicative of heterokaryotic incompatibility in fungi. No barrages are seen among the S14 strains.

#### Incomplete penetrance of *Spok2*

To investigate the nature of the 3-spored asci, tetrad dissections were conducted with asci from crosses between the *Psk-S* strains Wa63 and Us5, and the naïve strain Wa46. If the 3-spored asci were the result of a 4-spored ascus in which one of the spores aborted, all three spores should be heteroallelic for *Spok2*. If the 3-spored asci are the result of incomplete penetrance of the killing factor, two spores should be homoallelic for *Spok2* while the other spore should have no copy of *Spok2*. Unfortunately, spores from the crosses had very low germination rates (1/15 for Wa63 x Wa46 and 1/12 for Us5 x Wa46) as compared to other crosses (generally close to 100% germination). The progeny from the successfully germinated spores were backcrossed to the parental strains and also allowed to self to infer their *Spok2* genotype. Crosses with the Wa63/Wa46 progeny revealed it to be homoallelic for *Spok2*, and crosses with the Us5/Wa46 progeny revealed it to have no copy of *Spok2*. Both of these observations are consistent with the hypothesis for incomplete penetrance of *Spok2*.

#### Strain T and the original reports of spore killing in *Podospora*

The strain T has featured prominently in a number of important publications on spore killing in *Podospora*. It was one of the two strains investigated in the original description of spore killing by ***Padieu and Bernet*** (***1967***) (translated and reinterpreted by ***Turner and Perkins*** (***1991***)), it was the strain in which *Spok1* was described ***Grognet et al.*** (***2014***), and it was part of an investigation of spore killing in German strains of *Podospora* (***Hamann and Osiewacz, 2004***). Our results clearly demonstrate that two strains labeled as T (T_G_ and T_D_ herein) are not only different strains, but are differnt species. The description of spore killing in ***Padieu and Bernet*** (***1967***) matches our observations of crosses between T_G_ and the *Psk-S* strain Wa63, including incomplete penetrance as implied by the presence of 3-spored asci. Thus, we believe T_G_ to be representative of the original T strain. In light of this, we reinterpret the results of both ***Padieu and Bernet*** (***1967***) and ***Hamann and Osiewacz*** (***2004***) as per the interactions of the *Spok* genes.

In ***Padieu and Bernet*** (***1967***), they describe a cross between two strains: T and T’. They identify two genes (one present in T and the other in T’) which cause spore killing and interact as mutual killers. The gene from T has a killing percentage of 90%, while the one from T’ has a killing percentage of 40% and occasionally produces 3-spored asci. This fits well with a cross of *Psk-5* and *Psk-S* where *Psk-5* kills at 90% and *Spok2* of the *Psk-S* strain kills at 40%, but has incomplete penetrance resulting in 3-spored asci. Unfortunately strain T’ has to our knowledge not been maintained in any collections, so this cannot be confirmed experimentally. However, *Psk-S* strains are the most abundant phenotype from French, German, and Dutch populations (T’ was isolated in France along with T) (***van der Gaag et al., 2000***; ***Grognet et al., 2014***; ***Hamann and Osiewacz, 2004***).

In ***Hamann and Osiewacz*** (***2004***) they present a number of interesting observations. They report a new spore killer type, identify progeny that appear to demonstrate gene conversion of the killer locus, and observe apparent recombinant spore killer types. The study mostly centres around strain O, which they report to be of the same spore killer type as T_G_ and should thus be *Psk-5* given our results. As such, we suspect that their focal cross between O and Us5, a *Psk-S* strain, is the same as the Padieu and Bernet paper. We have independently confirmed that Us5 (kindly provided by A. Hamann and H. Osiewacz) is *Psk-S*, however strain O has not been maintained in any collection. They also state that strain He represents a new type of spore killer. However, with O classified as *Psk-5*, the interactions of He match that of a *Psk-1* strain. Furthermore, strain He exhibited no spore killing with a *Psk-1* strain from Wageningen. From the cross of O and Us5 they identify a number of progeny with unexpected genotypes. They interpret these genotypes as evidence for both gene conversion and recombinant spore killer types. However, under a two locus model of mutual killing, both effects can be explained by incomplete penetrance of *Spok2* (***Figure 2***). As the cross with Us5 showed a particularly high degree of anomalous results, it is possible that Us5 contains a unique allele of *Spok2* that is a particularly weak killer.

**Appendix 2 Figure 2.**
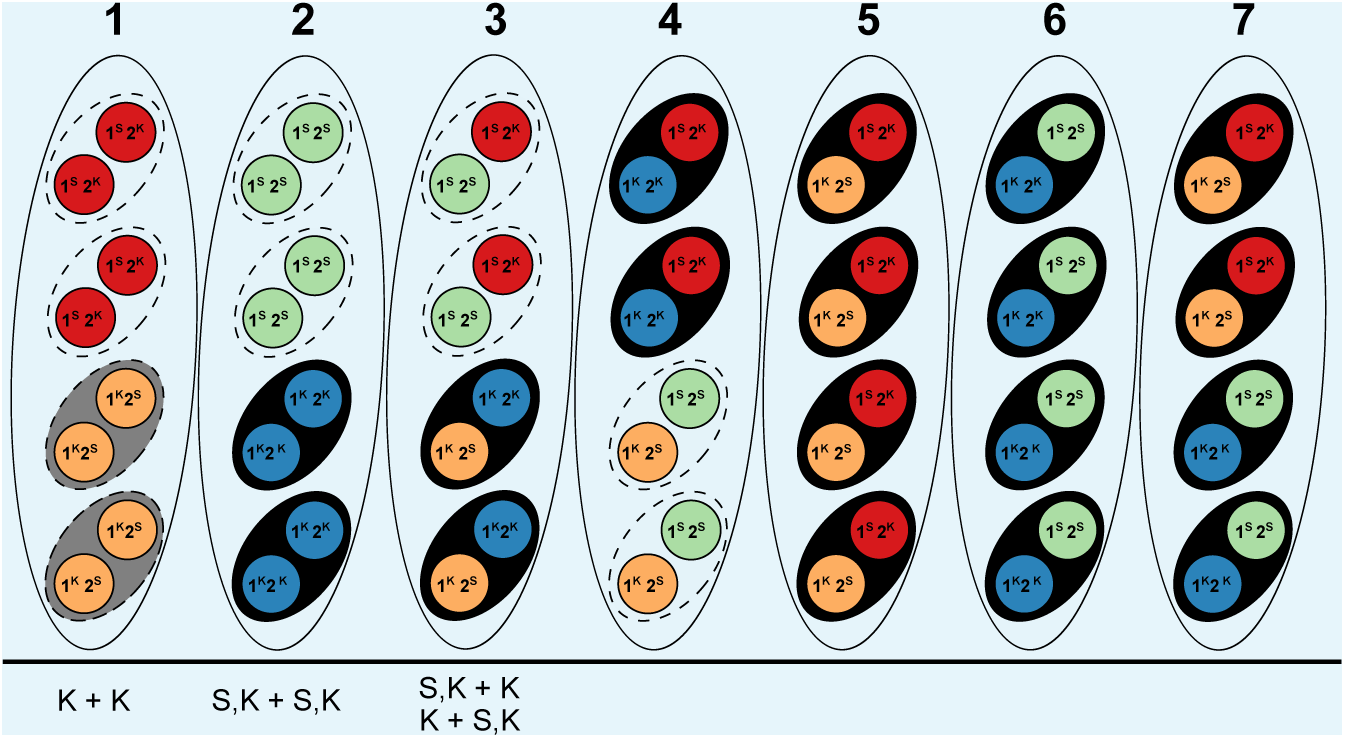
Explanation of results from ***Hamann and Osiewacz*** (***2004***) with information about *Spok* genes as described in the text. The seven asci represent the possible genotype combinations of a cross between a *Psk-5* strain and a *Psk-S* as illustrated in ***Turner and Perkins*** (***1991***). Black ovals represent the ascospores, dashed ovals represent killed spores, and coloured circles represent the individual nuclei, where each colour corresponds to a given genotype. Genotypes are annotated as per ***Turner and Perkins*** (***1991***), wherein locus 1 corresponds to a killer locus with 90% FDS, the *Psk-5 Spok* block, and locus 2 represents a killer locus with 40% FDS, *Spok2*. Red nuclei represent the *Psk-S* parental genotype with *Spok2*, orange nulcei represent the *Psk-5* parental genotype with *Spok3* and *Spok4*, green nuclei represent the recombinant genotype with no *Spok* genes, and blue nuclei represent the recombinant genotype with *Spok2, Spok3*, and *Spok4*. Note that *Spok3* and *Spok4* are linked and do not segregate independently. Below the asci are our interpretations of the annotations from ***Hamann and Osiewacz*** (***2004***). K + K strains would correspond to a strain with the *Psk-5* parental genotype of ascus type 1. These should experience mutual killing and produce empty asci, so the fact that they are observed from 4-spored asci suggests that when mutual killing occurs, 4-spores may be observed. However as no S + S strains were reported we can infer that only the *Psk-5* type (grey) may be viable. S,K+S,K strains are not indicative of a recombinant killer locus as suggested in the original work, but represent strains with all three *Spok* genes as produced in ascus type 2. The FDS frequencies reported suggest that the isolated strains are indicative of the blue nuclear genotype and not the green nuclear genotype. The S,K+K andK+ S,K strains are indistinguishable from each other and are indicative of the surviving spores of a type 3 ascus. These strains should exhibit spore killing when selfed due to the distribution of *Spok2*. Spore killing may not have been observed due to the incomplete penetrance of *Spok2*. In all cases, these strains should not have been isolated from 4-spored asci, indicating that either methodological issues occurred or that spore-killing may still produced 4-spored asci, but where the spores which should be absent are instead inviable.

