## Supplementary File Captions for "Combinations of *Spok* genes create multiple meiotic drivers in *Podospora*"

**Supplementary Figure Captions**

**Figure 1—Figure supplement 1.** Nucleotide alignment of the *Spok* homologs from the strains sequenced with long read technologies. The alignment includes the UTRs and intron. Start and stop codons are marked above the alignment track and the region of putative gene conversion are encased in red boxes.

**Figure 1—Figure supplement 2.** The expression of *Spok* genes based on RNAseq data. In order to ensure expression of the killer elements, we extracted RNA from an isolated self-killing heterozygote spore, produced by mating the monokaryotic *Psk* S_14_ backcrosses to a monokaryon S of opposite mating type (**A)**. The RNAseq data of one such culture (Psk7xS_14_ vs S) mapped to Wa58- (*Psk-7*) illustrates the expression of both *Spok3* (**B**) and *Spok4* (**C**). The expression of *Spok2* (**D**) and *Spok1* (**E**) in vegetative cells can be appreciated in the RNAseq data of Wa63- and T_D_+, respectively. In all cases, the predicted single exon of each *Spok* gene is shown bellow the reads, which in turn reveals an intron in the 5’UTR. Read colours follow the defaults in the genome browser IGV, with gray reads having typical expected insert size, and blue and red reads having slightly too small or too large insert size, respectively. The distributions above the reads correspond to the depth of coverage. Some sites appear polymorphic (blue and red bars in the depth distributions) due to low frequency mismapping of reads from other *Spok* homologs.

**Figure 1—source data 1.** Nucleotide alignment of *Spok* genes.

**Figure 1—source data 2.** Splits tree input file in Nexus format.

**Figure 2—Figure supplement 1.** Alignment of the *Spok* block from the *Psk-1* and *Psk-5* strains shows high overall collinearity. Colours are like in the **Figure 3** of the main text, except that the purple block highlights a large duplicate region unique to the T_G_ strain.

**Figure 2—Figure supplement 2.** Dot plot comparing the Wa87- *Spok* block (between and excluding genes Pa_3_945 and Pa_3_950) to the region containing *SpokΨ1* in Wa87- (Pa_5_10570 and Pa_5_10565). Red lines indicate collinear regions, while blue lines indicate inverted regions. The alignment was produced with NUCmer using the options -b 2000 -c 20 -p -maxmatch. Excluding the *Spok* homologs, there is little similarity between the two regions.

**Figure 2—Figure supplement 3.** Chromosomal segments remaining in the genomes after backcrossing of spore killer strains into the S background, along with the parental strains. Chromosomes were divided in 20 Kbp bins and coloured based on the count of SNPs called against the reference (Podan2). Bins identical to the reference (no SNPs) are not coloured. Below each chromosomal panel a cartoon in gray shows the length of the chromosome and the position of fixed genomic features present in all strains, including the centromere, the mating type (MAT) and *Spok2*. The centromere positions were approximated based on the genetic map of *P. anserina* (Espagne et al. 2008) and a large drop in GC content (not shown). When present, the positions of *Spok3* and *Spok4* were marked based on the insertion point of the *Spok* block in the long-read assemblies. Known *het* genes are marked and alleles of *het-s* and *het-c* are indicated in different colours.

**Figure 2—source data 1.** Fasta file of the *Spok* block from all strains.

**Figure 2—source data 2.** Fasta file of the *SpokΨ1* region from all strains.

**Figure 3—source data 1.** Fasta file of the *Spok2* region from all strains.

**Figure 3—source data 2.** Annotation file for TEs surrounding *Spok2*.

**Figure 4—Figure supplement 1.** Simplified killing hierarchy of the *P. anserina* spore killer types described in van der Gaag (2000). Killer capacity goes from maximum on top to null at the bottom. Red arrows signal killing dominance while blue lines mean mutual resistance, with corresponding killing percentages on the side. For example, according to van der Gaag 2000, *Psk-7* is mutually resistant with *Psk-1* (23% killing due to independent segregation), and it is dominant over *Psk-5* and all the other killer types. Notice that in the original work, the sensitive strains where in fact carriers of the *Spok2* gene, which is nearly cryptic due to its high frequency in the population. The killer types *Psk-6* and *Psk-3* are ignored (see main text).

**Figure 4—Figure supplement 2.** Genetic manipulations of *Spok* genes in the s strain background. For each cross, the *Spok*-genotype of the parents is given above the image of the rosettes. The s x s control cross yields 4-spored asci (**A**) and s x ΔSpok2 cross about 40% 2-spored asci (**B**). The *Spok3::PaPKS1* Δ*Spok2*} x ΔSpok2} and *Spok4::PaPKS1 ΔSpok2* x ΔSpok2 crosses yield 2-spored asci with two white spores (**C**, **D**). The *Spok3::PaPKS1d ΔSpok2* x ΔSpok2 shows 2-spored asci with two black spores and the *Spok3::PaPKS1 ΔSpok2* x *Spok3::PaPKS1d ΔSpok2* cross shows 4-spored asci with two white and two black spores as expected if the integration of *Spok3* downstream of the *PaPKS1* locus was successful (**E**, **F**). The Spok4::PaPKS1 ΔSpok2 x *Spok3::PaPKS1d ΔSpok2* cross is barren with close to 100 % empty asci (**G**). Crosses of *Spok3::PaPKS1* x s and *Spok4::PaPKS1* x s both result in 2-spored asci with two white spores (**H**, **I**). The *Spok3::PaPKS1* x *Spok3::PaPKS1 ΔSpok2* yields both 4-spored asci with two black and two white spores and 2-spored asci with black spores as expected if *Spok2* is inducing spore killing in this cross (**J**). The *Spok4::PaPKS1 ΔSpok2* x *Spok4::PaPKS1* cross is of poor quality due to the homogyzous deletion of *PaPKS1* but shows presence of some two-spored asci (marked with *) suggesting *Spok2* killing occurs. The *Spok3i::PaPKS1* x s (**L)** is identical to the *Spok3::PaPKS1* x s (**H**) suggesting no role for the intron in the spore killing action. The *Spok3* K204A was inserted at the *PaPKS1* locus and the resulting strain was crossed either with a *ΔSpok2* strain (**M**) or a *Spok3::PaPKS1d* (**N**). In both cases, 4-spored asci are produced, with two white and two black spores indicating that the mutant allele has lost spore killing function but retains resistance to *Spok3* killing.

**Figure 4—Figure supplement 3.** Plot comparing pooled sequencing data of progeny of 2-spored asci (left, n = 21) to progeny of 4-spored asci (right, n = 63) from a cross of *Psk-1* (Wa87+) and *Psk-5* (Y-). The Y-axis shows the frequency of each parental allele in the pooled samples, Wa87 (red) and Y (teal). In the 2-spored sample the region encompassing *Spok2* on Chromosome 5 is only represented by Wa87 alleles. This is the only region in the 2-spored pool with a complete skew and linkage to the centromere, as expected for a spore killing locus. Other skews from 50:50 ratios may be due to *het* genes, which can result in the death of nuclei when incompatible alleles are present in the same mycelium. Known *het* genes are marked. Skews specific to the 4-spore asci might be result of sibling competition effects during germination, imposed by the experimental design in the 4-spore asci (having two genotypes), but not in the 2-spore asci (only one genotype).

**Figure 4—source data 1.** Table with killing percentages for all crosses tested between strains.

**Figure 4—source data 2.** Table with killing percentages for test crosses to determine epistatic interactions.

**Figure 5—Figure supplement 1.** Amino acid alignment of the SPOK proteins from strains sequenced with long-read technologies. The N-terminus after the stop codon in *Spok2* and *Spok1* is not aligned due to a frame shift mutation (see nucleotide alignment). Domains and conserved motifs from Figure 5 are indicated in matching colours. Point mutations investigated in this study are marked with yellow triangles. The C-terminus truncation of SPOK3 is marked with a blue dashed line, while the two putative NLS signals are shaded in blue at the top of the alignment. Note that the *Spok3(1-490)* construct also has a putative NLS.

**Figure 5—Figure supplement 2.** Predicted nuclease domain of SPOK proteins. The alignment of the SPOK3 domain 2 and the nuclease domain of the HsdR restriction enzyme from *Vibrio vulnificus* generated by HHPred is given (**A**). The figure gives the secondary structure prediction for the query and target, labelled ss (pred), as well as a consensus sequence for the query and target, labelled cons. as the actual secondary structure of the target in dssp code. The catalytic core residues of the PD-(D/E)XK are underlined as well as the QXXXY motif important for nuclease activity. In addition, the positions mutated in the SPOK2 in strain A are given. The arrow points to the position of the two amino acid insertion in that strain. A structural model of domain 2 of SPOK3 based on a contact map generated with RaptorX is given (**B**). Positions of the catalytic core residues are given with the same colour code as in (**A**). Again the positions mutated in strain A are shown. A short alignment of the wild-type SPOK2 and mutated SPOK2 sequence as found in strain A is shown. (**C**) The structure of the HsdR nuclease domain (pdb 3H1T) in which the homologous positions have been highlighted according to the alignment given in (**A**). Note the close spatial proximity of the catalytic lysine and the site of the two amino acid insertion in strain A in both cases.

**Figure 5—source data 1.** Amino acid alignment of the SPOK proteins in the *Podospora* complex.

**Figure 5—source data 2.** Gremlin amino acid alignment of SPOK proteins closely related to those in *Podospora*.

**Figure 5—source data 3.** Transformation efficiency of *Spok3* manipulations.

**Figure 6—source data 1.** Codon-guided alignment of homologs closely related to the *Podospora* SPOKs.

**Supplementary file 1.** Statistics from PacBio and Nanopore assemblies.

**Supplementary file 2.** Statistics from SPAdes assemblies.
