## Supplementary material for "Combinations of *Spok* genes create multiple meiotic drivers in *Podospora*": Figure 5--source data 3

**Transformation efficiency of *Spok3* D667A with or without wild-type *Spok3***

| vector 1 (with selection marker) | vector 2  (for rescue) | transfo. #1 | transfo. #2 | total | % of control |
| --- | --- | --- | --- | --- | --- |
| *Spok3* | control | 160 | 55 | 215 | - |
| *Spok3* D667A | control | 23 | 4 | 27 | 12.5% |
| *Spok3* | *Spok3* | 140 | 42 | 182 | 84.6% |
| *Spok3* D667A | *Spok3* | 134 | 72 | 206 | 95.8% |

**Transformation efficiency of *Spok3* C493A C497A, C511A and C511S with or without wild-type *Spok3***

| vector 1 (with  selection marker) | vector 2  (for rescue) | transfo. #1 | transfo. #2 | total | % of control |
| --- | --- | --- | --- | --- | --- |
| *Spok3* | control | 80 | 75 | 155 | - |
| *Spok3* C493A C497A | control | 15 | 8 | 23 | 14,8% |
| *Spok3* C511S | control | 5 | 15 | 20 | 12.9% |
| *Spok3* C511A | control | 5 | 8 | 13 | 8.3% |
| *Spok3* C493A C497A | *Spok3* | 76 | 56 | 132 | 85.1% |
| *Spok3* C511S | *Spok3* | 55 | 57 | 112 | 72.2% |
| *Spok3* C511A | *Spok3* | 67 | 51 | 118 | 76.1% |

**Transformation efficiency of *Spok3* (1-490) with or without wild-type *Spok3***

| vector 1 | vector 2  (for rescue and with selection marker) | transfo. #1 | transfo. #2 | total | % of control |
| --- | --- | --- | --- | --- | --- |
| control | *Spok3* | 315 | 351 | 666 | - |
| *Spok3* (1-490) | control | 7 | 7 | 14 | 2.1% |
| *Spok3* (1-490) | *Spok3* | 262 | 299 | 563 | 84.5% |
