## Supplementary figures and images for "Combinations of *Spok* genes create multiple meiotic drivers in *Podospora*"

### Figure 1--Figure supplement 1

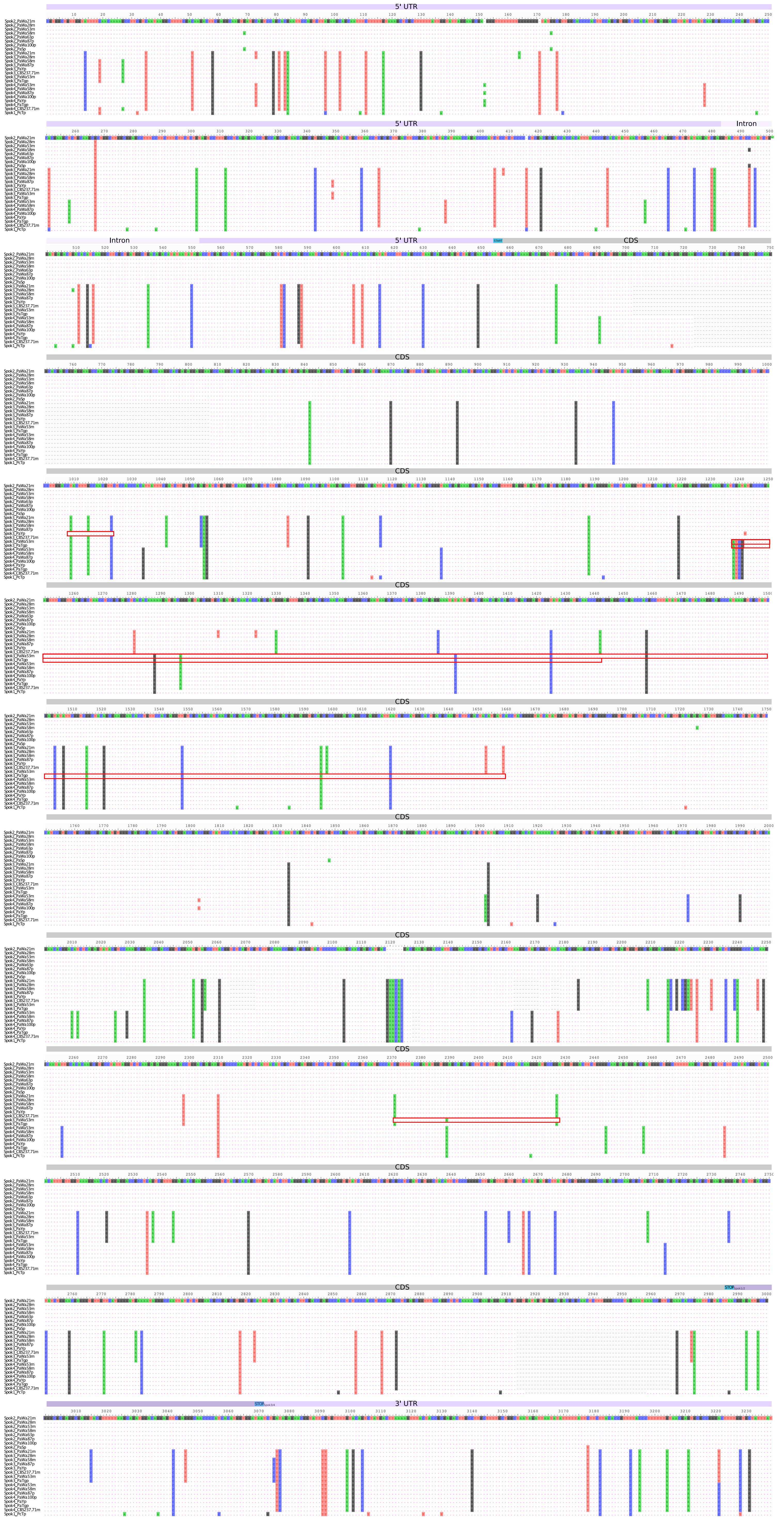

### Figure 1--Figure supplement 2

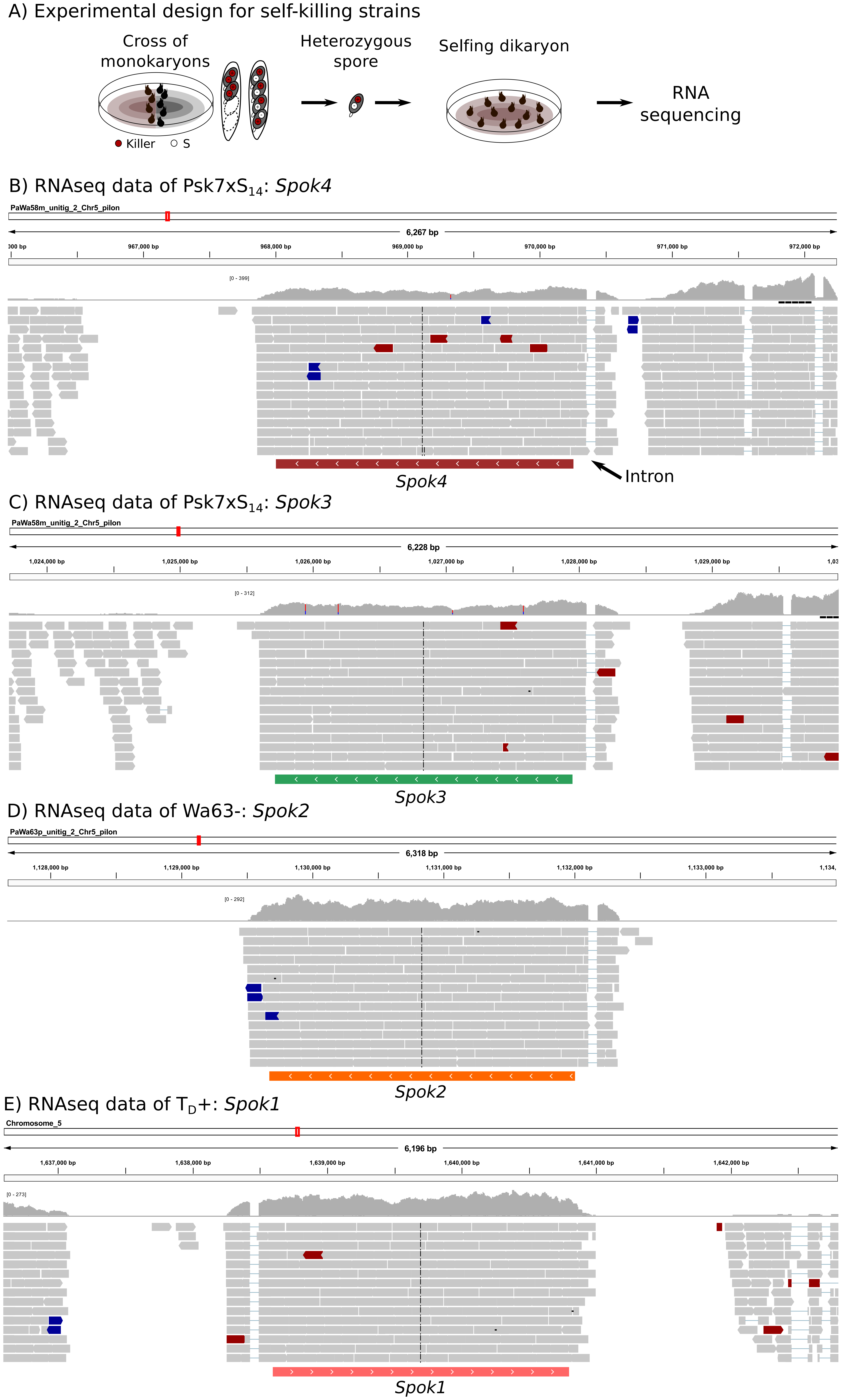

### Figure 2--Figure supplement 1

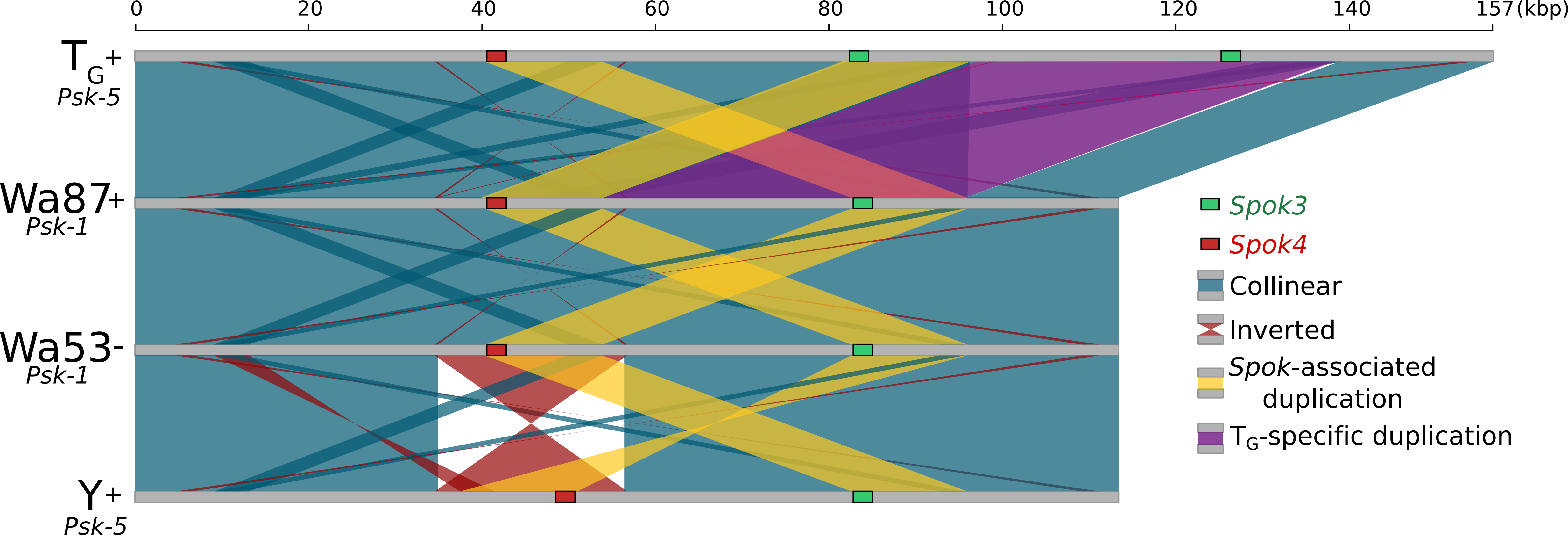

### Figure 2--Figure supplement 2

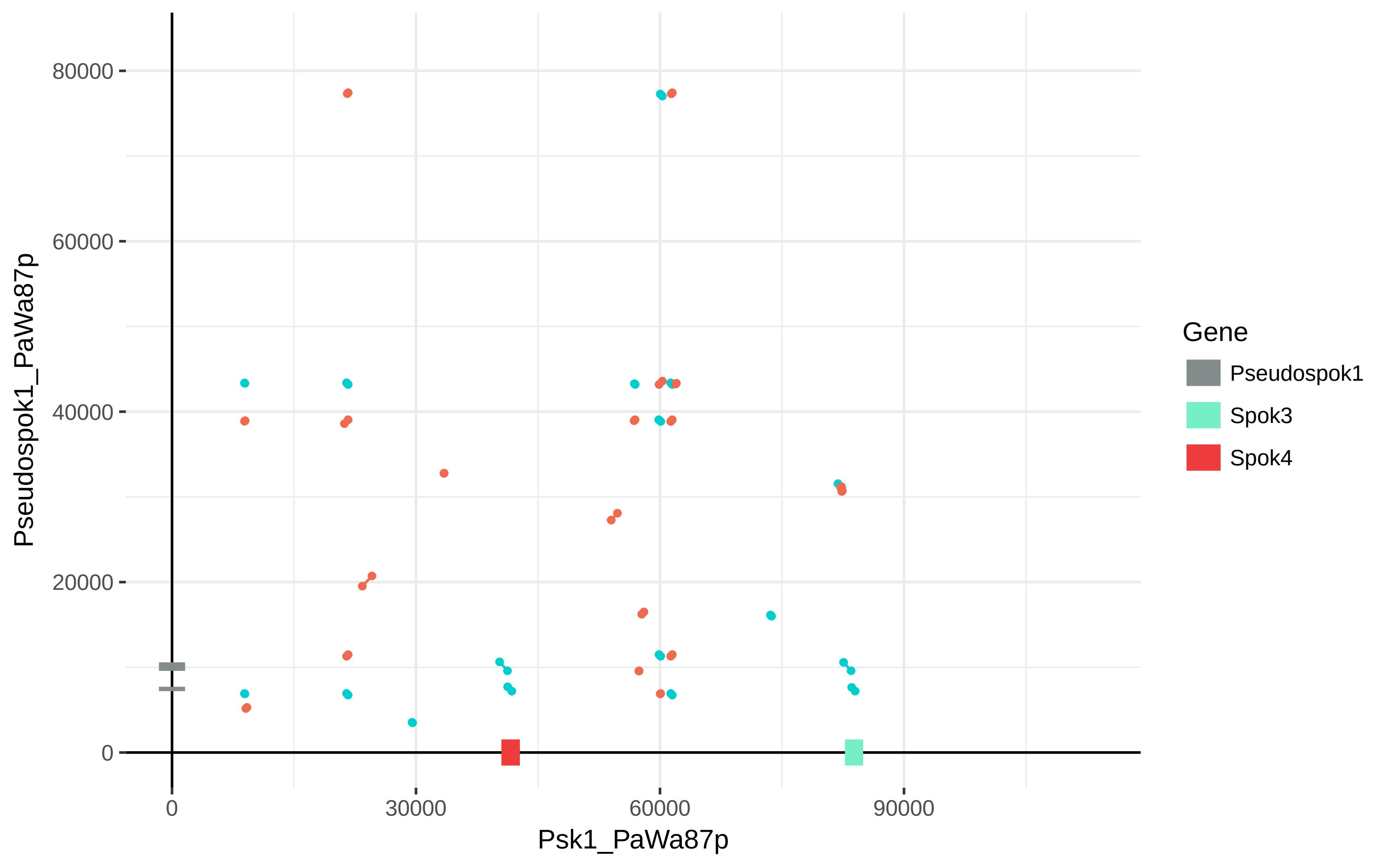

### Figure 2--Figure supplement 3

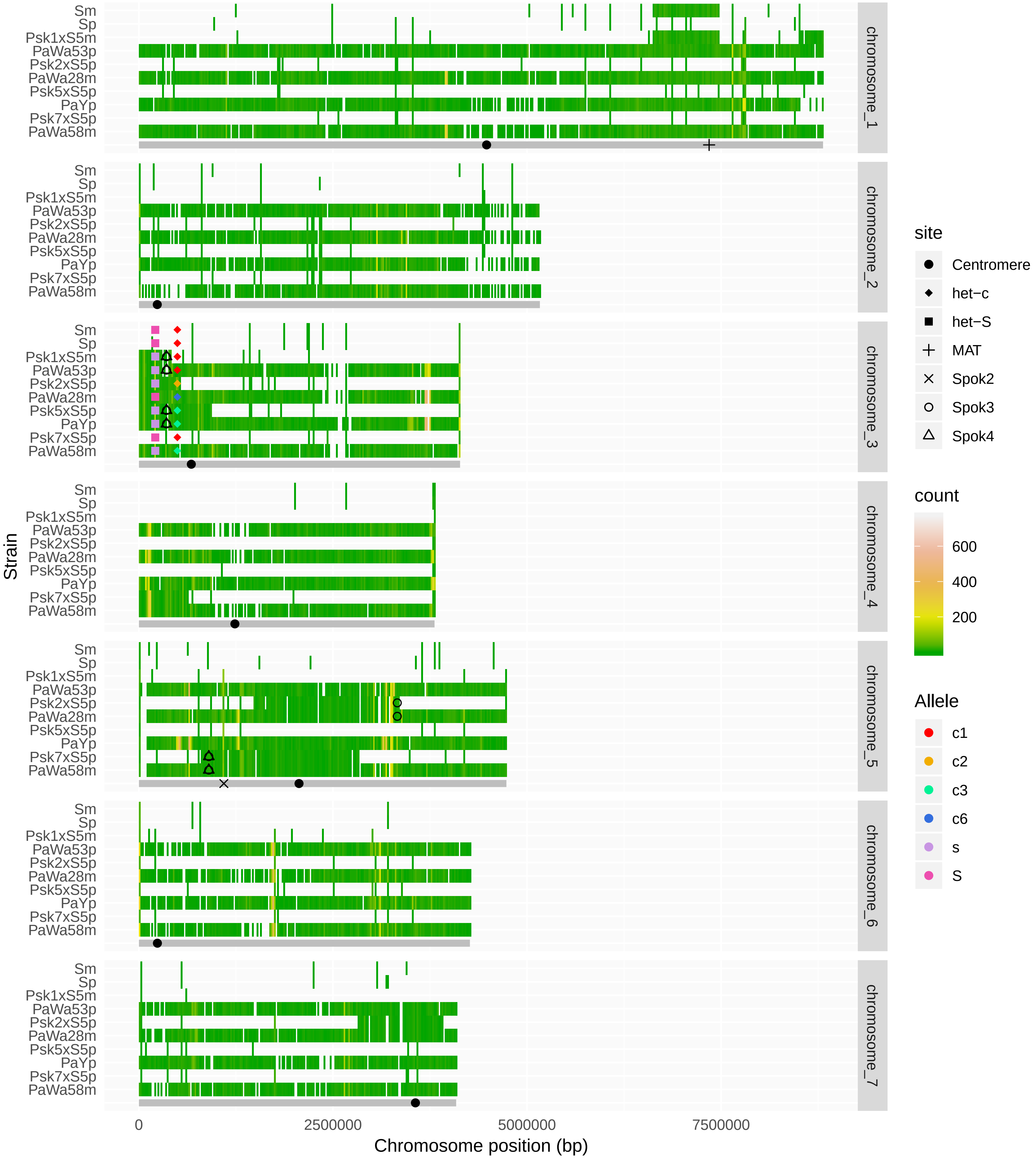

### Figure 4--Figure supplement 1

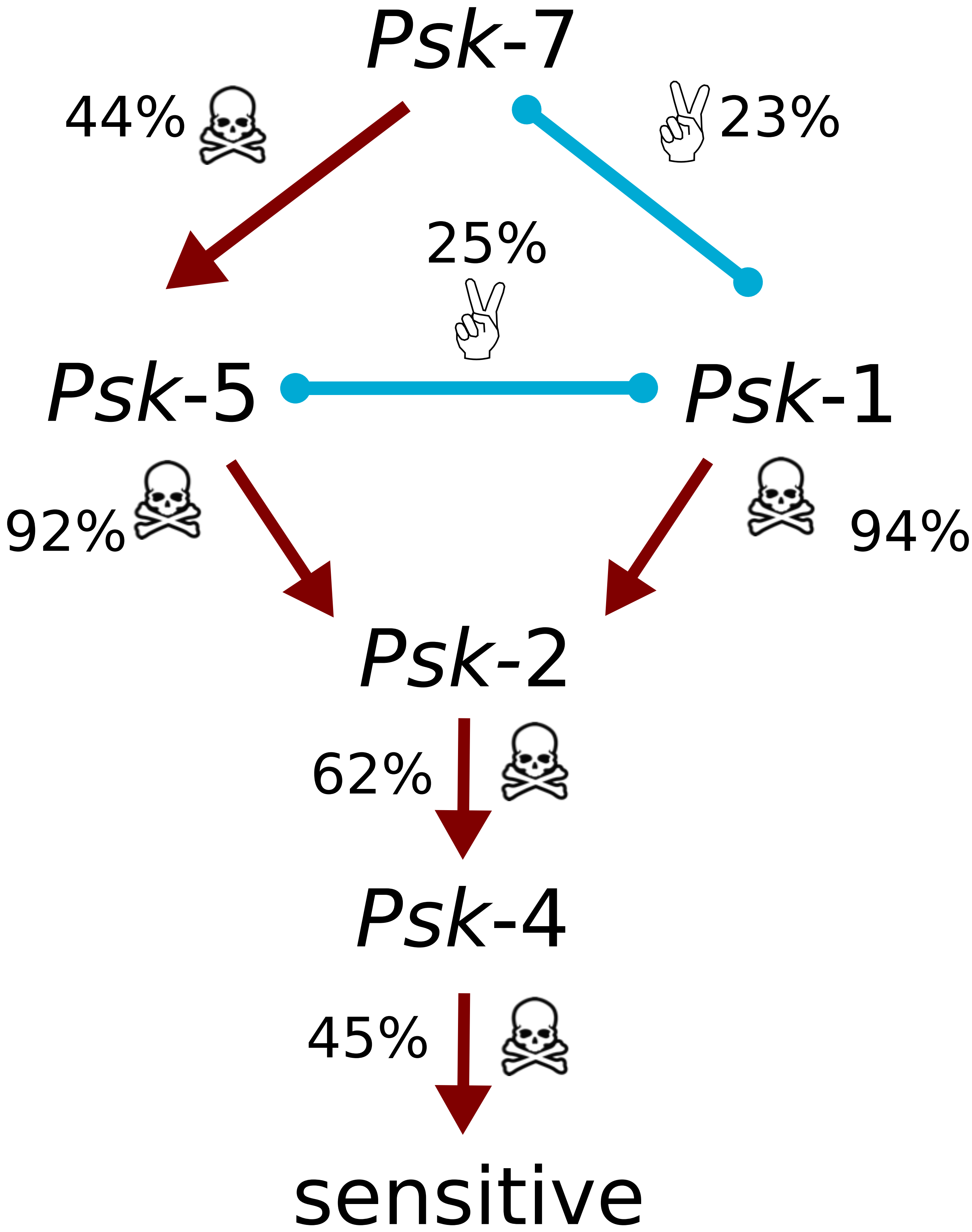

### Figure 4--Figure supplement 2

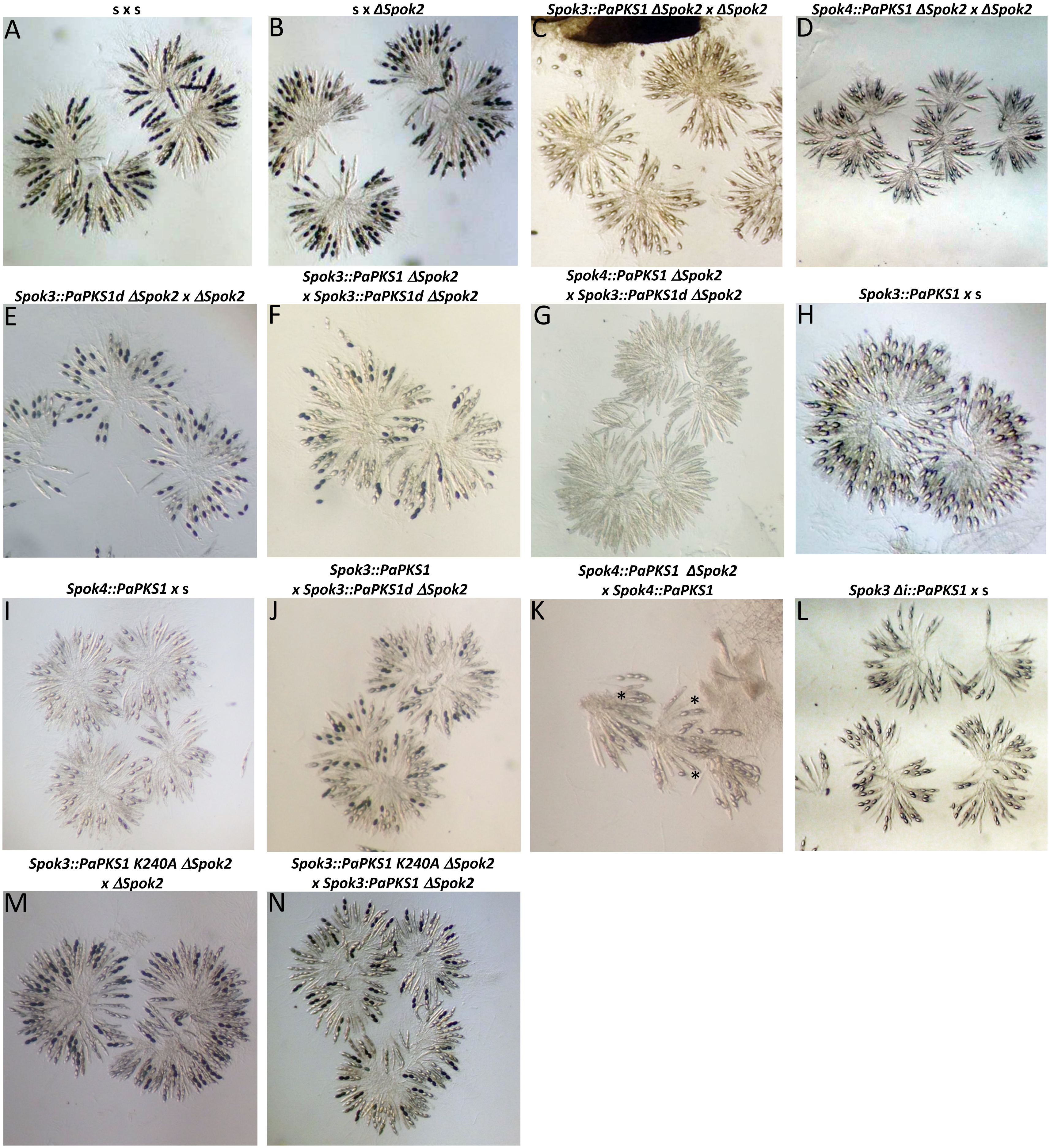

### Figure 4--Figure supplement 3

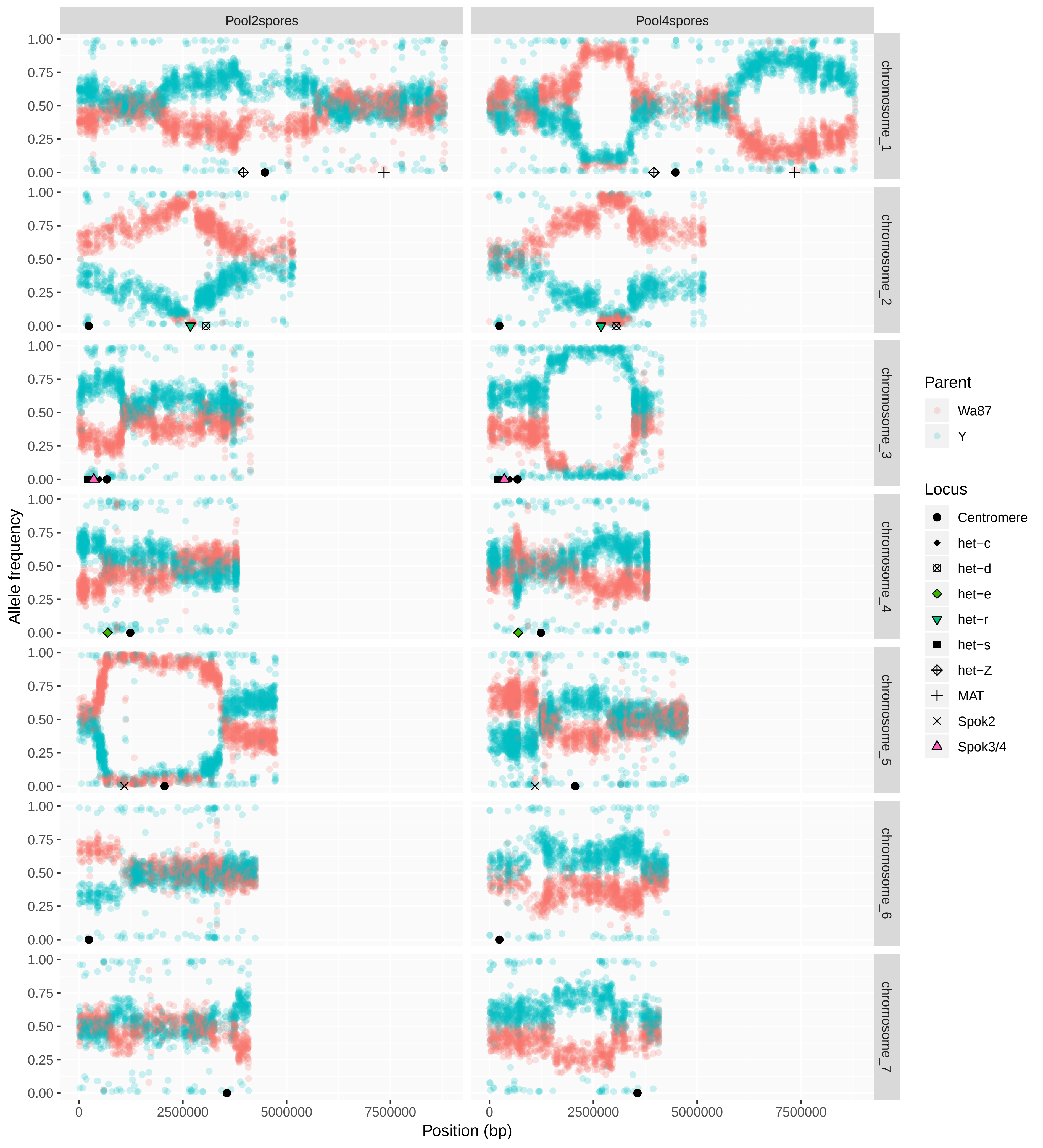

### Figure 5--Figure supplement 1

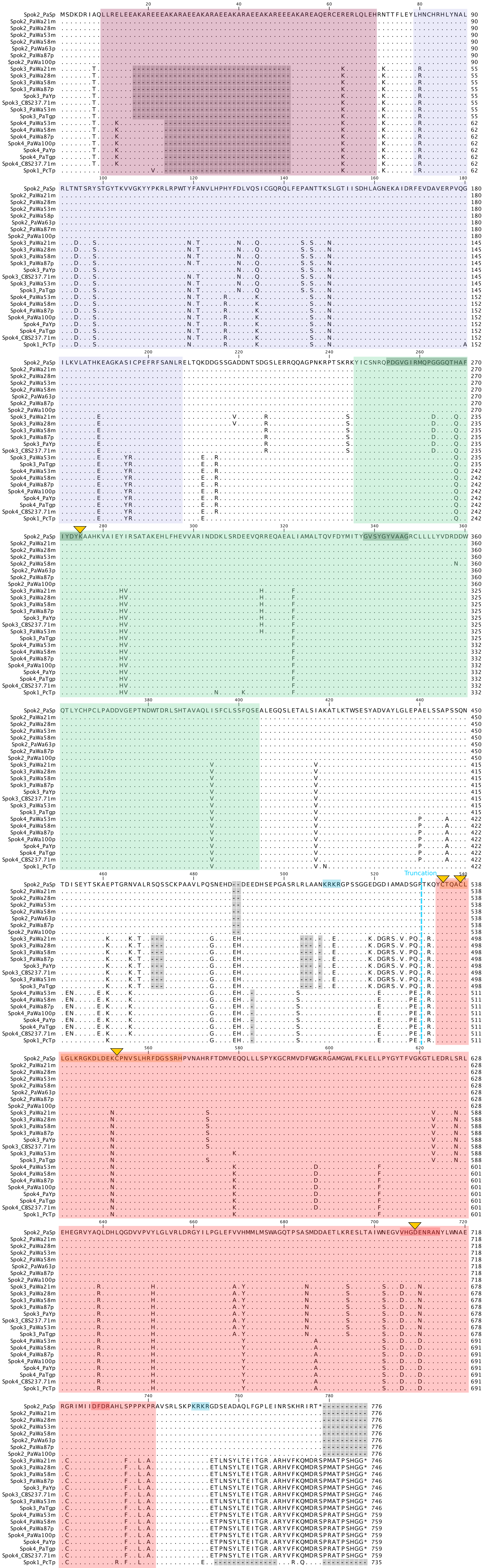

### Figure 5--Figure supplement 2

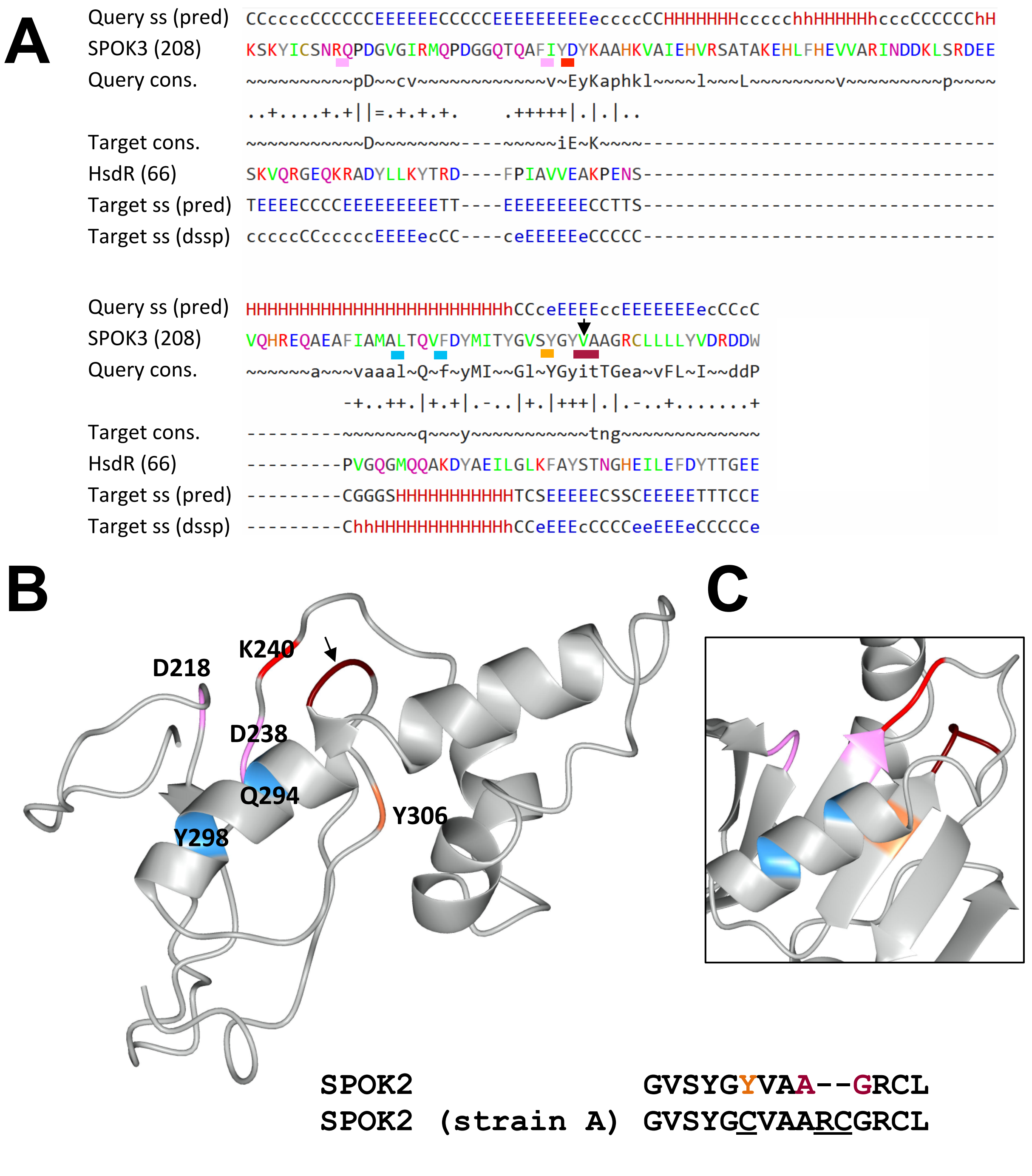
